# An artificial self-assembling nanocompartment for organising metabolic pathways in yeast

**DOI:** 10.1101/2021.01.30.428974

**Authors:** Li Chen Cheah, Terra Stark, Lachlan S. R. Adamson, Rufika S. Abidin, Yu Heng Lau, Frank Sainsbury, Claudia E. Vickers

## Abstract

Metabolic pathways are commonly organised by sequestration into discrete cellular compartments. Compartments prevent unfavourable interactions with other pathways and provide local environments conducive to the activity of encapsulated enzymes. Such compartments are also useful synthetic biology tools for examining enzyme/pathway behaviour and for metabolic engineering. Here, we expand the intracellular compartmentalisation toolbox for budding yeast (*Saccharomyces cerevisiae*) with engineered Murine polyomavirus virus-like particles (MPyV VLPs). The MPyV system has two components: VP1 which self-assembles into the compartment shell; and a short anchor, VP2C, which mediates cargo protein encapsulation via binding to the inner surface of the VP1 shell. Destabilised GFP fused to VP2C was specifically sorted into VLPs and thereby protected from host-mediated degradation. In order to access metabolites of native and engineered yeast metabolism, VLP-based nanocompartments were directed to assemble in the cytosol by removal of the VP1 nuclear localisation signal. To demonstrate their ability to function as a metabolic compartment, MPyV VLPs were used to encapsulate myo-inositol oxygenase (MIOX), an unstable and rate-limiting enzyme in D-glucaric acid biosynthesis. Strains with encapsulated MIOX produced ~20% more D-glucaric acid compared to controls expressing ‘free’ MIOX - despite accumulating dramatically less expressed protein - and also grew to higher cell densities. These effects were linked to enzyme stabilisation and mitigation of cellular toxicity by the engineered compartment. This is the first demonstration in yeast of an artificial biocatalytic compartment that can participate in a metabolic pathway and establishes the MPyV platform as a promising synthetic biology tool for yeast engineering.

## INTRODUCTION

Intracellular metabolic compartments are ubiquitious in nature. Common examples are the membrane-bound organelles such as the mitochondrion, chloroplast, and peroxisome. In addition, certain prokaryotes express protein-based metabolic compartments such as bacterial microcompartments (BMCs) and encapsulins. Compartments create discrete favourable environments for otherwise incompatible reactions, while minimising unproductive interactions that lead to toxicity and metabolite loss^1,2^. Other functions of compartments include spatially organising successive enzymes in a pathway to improve pathway efficiency and increasing the local substrate concentration to favour a particular reaction^1,2^. In some cases, engineered compartmentalisation has been found to impart useful properties on enzymes such as improved activity and stability^3,4^.

The bottom-up reconstruction of ‘synthetic organelles’ has recently been explored using self-assembling protein compartments^4,5^. These compartments can be used to encapsulate metabolic enzymes in an engineered pathway to enhance chemical bioproduction. The use of heterologous compartments reduces undesirable cross-talk with host cell metabolism and potentially enables finer control of the reaction environment. Furthermore, the inherent programmability of protein-based compartments means their permeability and surface chemistry can be tuned to favour a particular reaction. For instance, the pore size and charge may be engineered to favour the influx of substrates and/or minimise the efflux of intermediate metabolites^6,7^. As each type of protein compartment has characteristics that may make it better suited to different applications, it is useful to continually explore and develop new compartment platforms. One property that is particularly relevant for biocatalysis is its permeability to substrates – highly porous compartments such as the bacteriophage P22 procapsid permit free diffusion of small molecules^8^, while compartments with small pores such as bacterial microcompartments and encapsulins allow selective metabolite exchange^9,10^. Harnessing the natural diversity of self-assembling compartment structures thus allows us to generate a suite of tools that can fulfil distinct niches.

Here, we present an artificial metabolic nanocompartment for budding yeast (*Saccharomyces cerevisiae*) based on the Murine polyomavirus virus-like particle (MPyV VLP). MPyV coat proteins are known to self-assemble in various heterologous eukaryotic expression hosts, including yeasts^11–13^. The MPyV VLP makes an attractive base for constructing designer compartments due to its amenability to engineering and ability to selectively package cargo proteins^14,15^. The MPyV shell is porous, with gaps between capsomeres (virus assembly subunits) as well as a central 8.6 Å pore through each capsomere^16,17^; this could potentially enable access of encapsulated enzymes to small molecule substrates. The VLP exterior can be functionalised with various domains by insertion into loop regions, a property which has previously been exploited for modular antigen display^18–20^. The compartment is ~50 nm in diameter and has a theoretical maximum loading of 72 cargo proteins per particle, providing a larger capacity than a previously reported artificial nanocompartment system for yeast^21^. By coat protein engineering, we developed an orthogonal compartment that localises to the yeast cytoplasm and packages an exceptionally high density of cargo proteins. The MPyV platform was then applied towards the *in vivo* stabilisation of a metabolic enzyme, which resulted in improved product titres as well as increased cell growth. The MPyV platform provides novel capabilities, expanding the *in vivo* protein scaffolding and compartmentalisation toolbox for this important bioproduction chassis.

## RESULTS AND DISCUSSION

### Design and characterisation of a synthetic MPyV-based yeast nanocompartment

The engineered MPyV system has two protein components: VP1, which forms the compartment shell; and VP2C, a short anchor for directing cargo protein encapsulation^13,22^ (Figure 1a). VP1 assembles into pentamers, which then further self-assemble into a VLP nominally composed of 72 pentamers (360 VP1 monomers). Each VP1 pentamer can bind one VP2C anchor and, by extension, a cargo protein translationally fused to VP2C. Exploiting the VP2C-VP1 interaction allows specific packaging of the cargo protein of interest during assembly of the VLP. Since VP2C is not essential for pentamer or VLP formation, the assembled particles may contain variable numbers of ‘empty’ pentamers, as depicted in Figure 1a.

**Figure 1.**
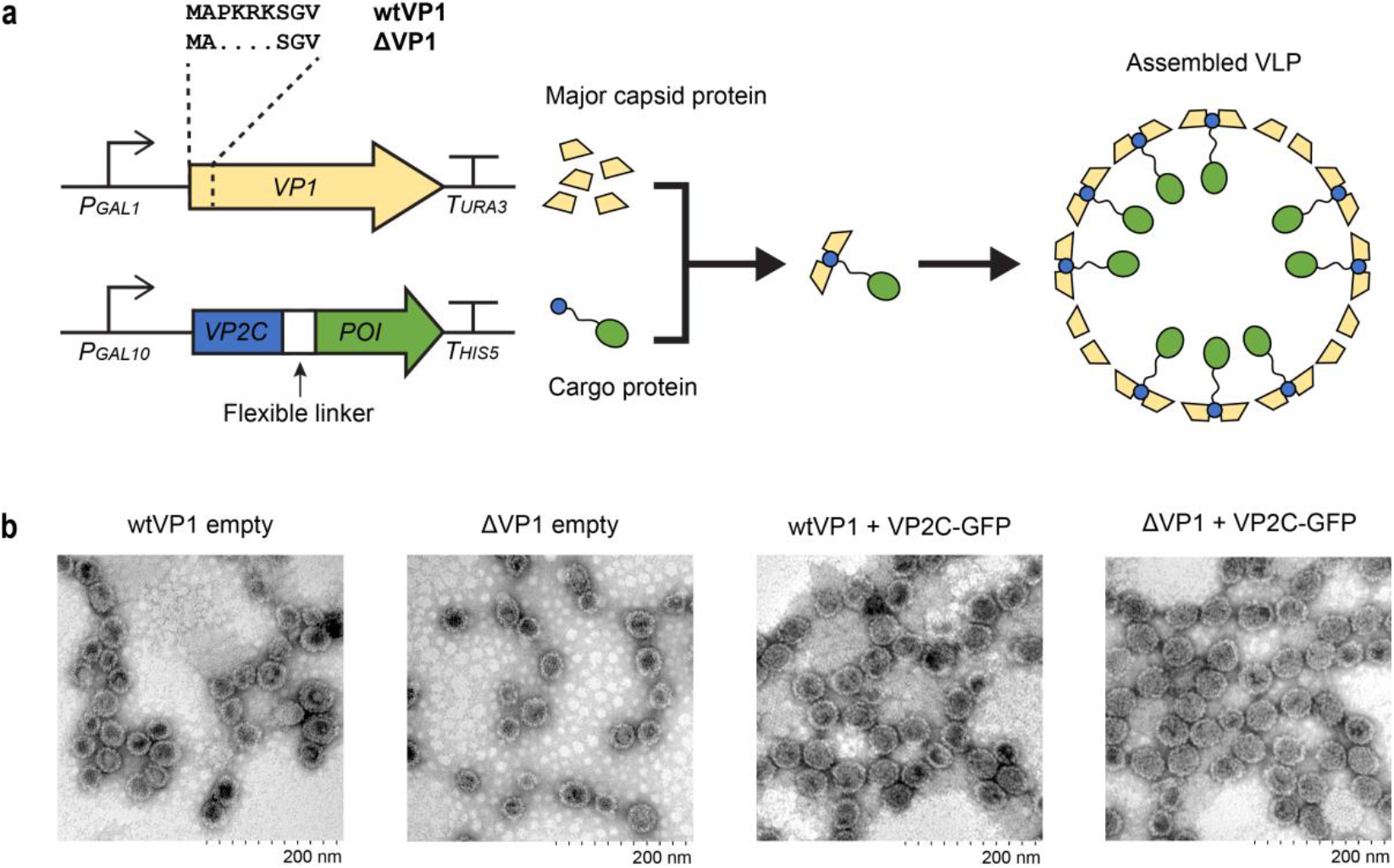
The MPyV nanocompartment platform for yeast. (a) MPyV virus-like particles (VLPs) are formed by the self-assembly of two protein components, VP1 (wt or an NLS-deletion mutant, Δ) and VP2C linked to the cargo protein of interest (‘POI’). (b) Transmission electron micrographs of purified VLPs expressed in the absence and presence VP2C-GFP.

When expressed in yeast, wild-type MPyV VP1 (wtVP1) forms VLPs in the nucleus^11^. For this project, we were interested in designing a cytoplasmic compartment system because of the larger diversity of metabolic pathways and processes in the yeast cytoplasm compared to the nucleus. In our previous work on plant-expressed MPyV VLPs^13^, deletion of a putative nuclear localisation signal on VP1 (mutant to be referred to as ‘ΔVP1’ hereafter; Figure 1a) abolished exclusive nuclear localisation while maintaining VLP assembly capabilities. We sought to assess the suitability of ΔVP1 for generating yeast compartments, comparing localisation, assembly and capacity for cargo encapsulation with wtVP1 by first testing the system using yeast-enhanced GFP^23^ as the model cargo protein.

ΔVP1 and wtVP1 were expressed either alone (forming empty VLPs) or coexpressed with VP2C-GFP (forming GFP-loaded VLPs) using strong galactose-inducible promoters (Figure 1a). Purified ΔVP1 and wtVP1 VLPs share similar morphology under transmission electron microscopy (TEM) (Figure 1b). Expression of VP2C-GFP led to effective cargo packaging in both VLPs as indicated by SDS-PAGE (Figure 2a) and native agarose gel electrophoresis (Figure 2b). The cargo loading was estimated to be ~59 GFP per wtVP1 VLP and ~72 GFP per ΔVP1 VLP by SDS-PAGE densitometry. Samples encapsulating GFP always exhibited two cargo bands in SDS-PAGE. The identity of both visible cargo bands was verified by anti-GFP western blot (Figure S1a) and N-terminal sequencing confirmed that the smaller cargo protein was a cleavage product of VP2C-GFP (Figure S1b). Using a pull-down assay in *E. coli*, we show that an even shorter truncation still allows binding to the VP1 pentamer (Figure S2), though it is not clear whether degradation of the N-terminus occurs prior to, or after, VP1 binding. Nevertheless, no other bands were seen on SDS-PAGE other than that of VP1 and VP2C-GFP, confirming the specificity of VP2C-directed cargo packaging. Removing the VP2C anchor from GFP led to VLPs without detectable GFP (Figure S3), indicating that the contribution of random, ‘statistical’ encapsulation is negligible.

**Figure 2.**
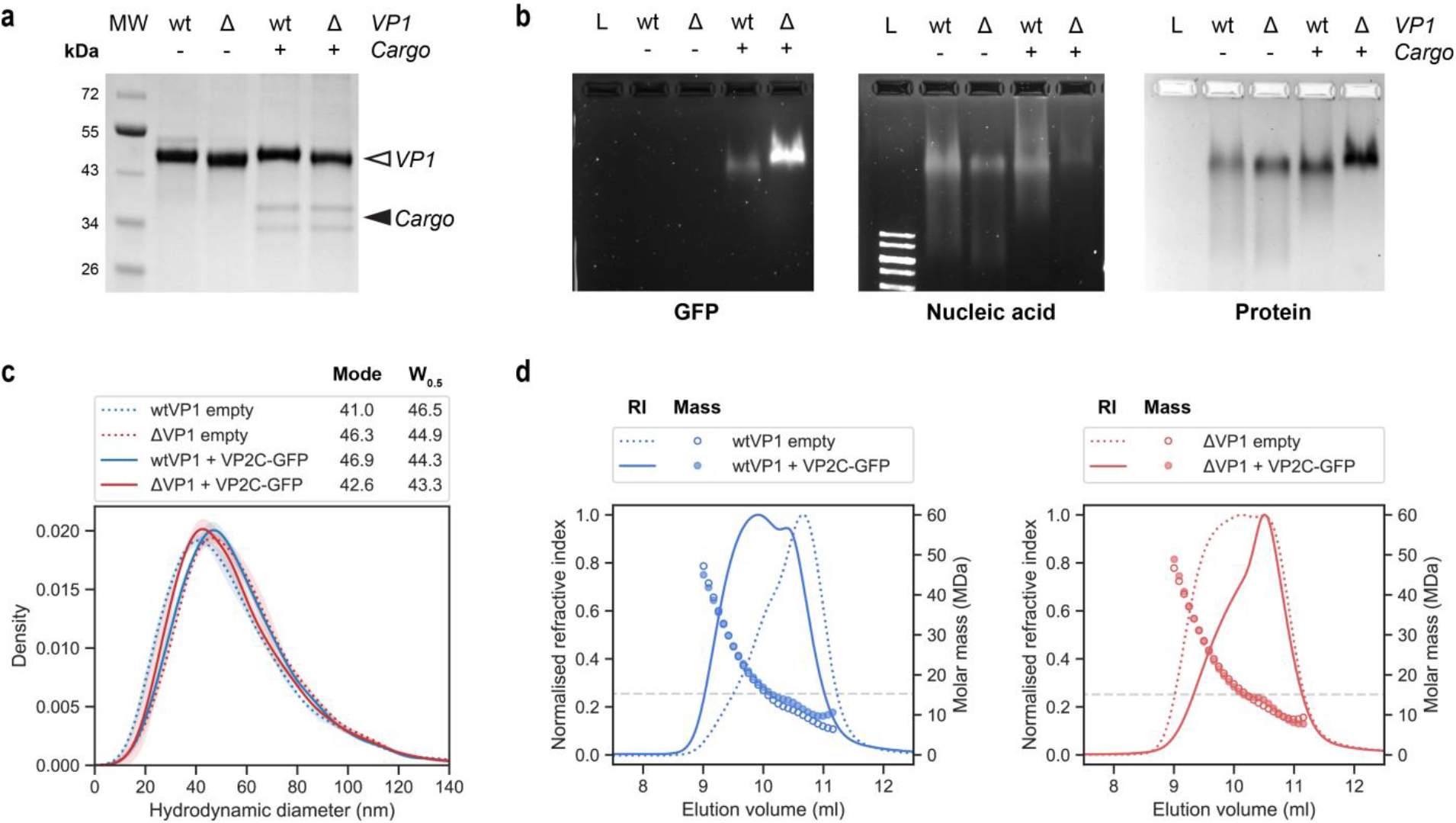
VLP characterisation. (a) SDS-PAGE gel of purified VLP samples stained with Coomassie blue. Arrows show the position of VP1 and cargo bands. ‘MW’ = protein molecular weight marker. (b) Native gel electrophoresis of purified particles. Samples (3 μg) were loaded on a 1% agarose gel alongside 0.5 μg of a DNA molecular ladder (lane L). GFP signal from intact particles can be visualised with blue light illumination and a 530 nm emission filter. Nucleic acid and protein were stained with GelRed and Coomassie blue respectively. (c) Particle size distributions, measured with nanoparticle tracking analysis (NTA). The mode and width at half height (W0.5) of each distribution is indicated. (d) Changes in size distribution and molar mass of wtVP1 and ΔVP1 VLPs with GFP loading, as determined by SEC-MALS. Refractive index, RI (normalised to the mode) is shown as lines and molar mass is shown as circles. The dashed light grey line indicates the theoretical mass of empty VLPs corresponding to each VP1 variant.

Empty MPyV VLPs have been reported to non-specifically encapsulate genomic and plasmid DNA when expressed in yeast^11^. Consistent with this, considerable nucleic acid staining for wtVP1 VLPs was observed on the native agarose gel (Figure 2b). Nucleic acid encapsulated in ΔVP1 VLPs was presumably RNA; however, it was greatly reduced compared to wtVP1, likely from deletion of a number of positively charged residues in the mutation^24,25^ and shifting of the site of assembly away from the nucleus^26,27^. Decreased nucleic acid encapsulation by ΔVP1 compared to wtVP1 has also been observed with plant-expressed MPyV VLPs^13^. Minimisation of nucleic acid encapsulation is desirable to maximise the effective capacity available for compartmentalisation of target proteins. The presence of encapsulated GFP also reduces nucleic acid capture for both wtVP1 and ΔVP1. The protruding VP2C-GFP is presumably sterically ‘blocking’ the lumen-facing surface of VP1 pentamers, reducing their availability for nucleic acid binding.

Nanoparticle tracking analysis (NTA) detected slight differences in the particle size distribution between compartment types (Figure 2c). NTA is a sensitive imaging method that calculates the size of individual particles in solution based on its Brownian motion^28^. For both VP1 variants, cargo encapsulation slightly narrows the width at half-height (W_0.5_) of the distribution, indicating an increase in size uniformity. Although the effect here is small, it echoes previous findings for MPyV VLPs assembled in insect cells where coexpression with full-length VP2 resulted in fewer aberrant-sized particles^29^. In an *in vitro* MPyV study, fluorescent protein encapsulation using the VP2C anchor led to a loading density-dependent reduction in W_0.5_^22^. However, heterologous cargo encapsulation *in vivo* had opposing effects on the average particle size for the VP1 variants; average particle size increased for wtVP1 and decreased for ΔVP1 (Figure 2c). When separated by size-exclusion chromatography (SEC), each VLP type exhibited complex elution profiles which appear to comprise of at least two overlapping peaks suggestive of particle sub-populations (Figure 2d). Comparing the refractive index profiles, cargo encapsulation by wtVP1 increased the proportion of larger sized VLPs (height of earlier sub-peak increases) while the opposite is true for ΔVP1. It is unclear why this might be, although the different biochemical environment of the inner surfaces of the two shell variants may result in varying interactions with the cargo. We do not expect the size variability to affect the performance of MPyV as a biocatalytic compartment.

Multi-angle light scattering (MALS) of SEC elutions showed that the molar mass of purified VLPs had a broad distribution that was centred around 15 MDa, the theoretical mass of a 72-pentamer assembly (Figure 2d). The ability of MPyV to form VLPs of different sizes has previously been observed *in vitro*^22,30^ and *in vivo*^13,31^. Interestingly, the mass of loaded and empty VLPs were similar up to ~10.2 ml, when GFP-loaded VLPs exhibit a pronounced mass bump. Although this remains to be investigated, it may indicate a bias in nucleic acid capture and cargo loading towards specific VLP sizes. For both shell variants, GFP loading appeared to result in a subpopulation of particles with a peak elution at ~10.5 ml (Figure 1d).

### Removing the VP1 nuclear localisation signal leads to cytoplasmic compartments and improved cargo capture

VP2C-GFP is expected to diffuse freely through the yeast nuclear pore complex^32^, making it a suitable reporter for the subcellular location of assembled compartments. To distinguish encapsulated cargo from excess ‘free’ cargo, we destabilised GFP by adding a C-terminal degradation signal from mouse ornithine decarboxylase^33^. The resulting high-turnover reporter, GFP_Deg_, has a half-life of ~10 min in yeast^33^. Similar strategies have previously been applied to study other protein compartments *in vivo*^21,34,35^ and to target competing enzymes for metabolic engineering^36–38^. Incubation of cultures with the protein synthesis inhibitor cycloheximide allows ‘clearing’ of unencapsulated cargo proteins, leaving only the signal from encapsulated GFP_Deg_ (Figure 3a). This was verified by separating VLP-associated GFP_Deg_ from free GFP_Deg_ by ultracentrifugation of whole cell lysates through an iodixanol cushion (Figure 3b). Even against an autofluorescent background, cycloheximide treatment led to a clear reduction in fluorescence signal in the upper fraction (free proteins), but not the dense VLP fraction. Fusion to the VP2C anchor was required for the sedimentation of GFP_Deg_ signal (Figure S3). The ΔVP1 strain also exhibited more GFP_Deg_ signal in the VLP fraction than the wtVP1 strain (Figure 3c).

**Figure 3.**
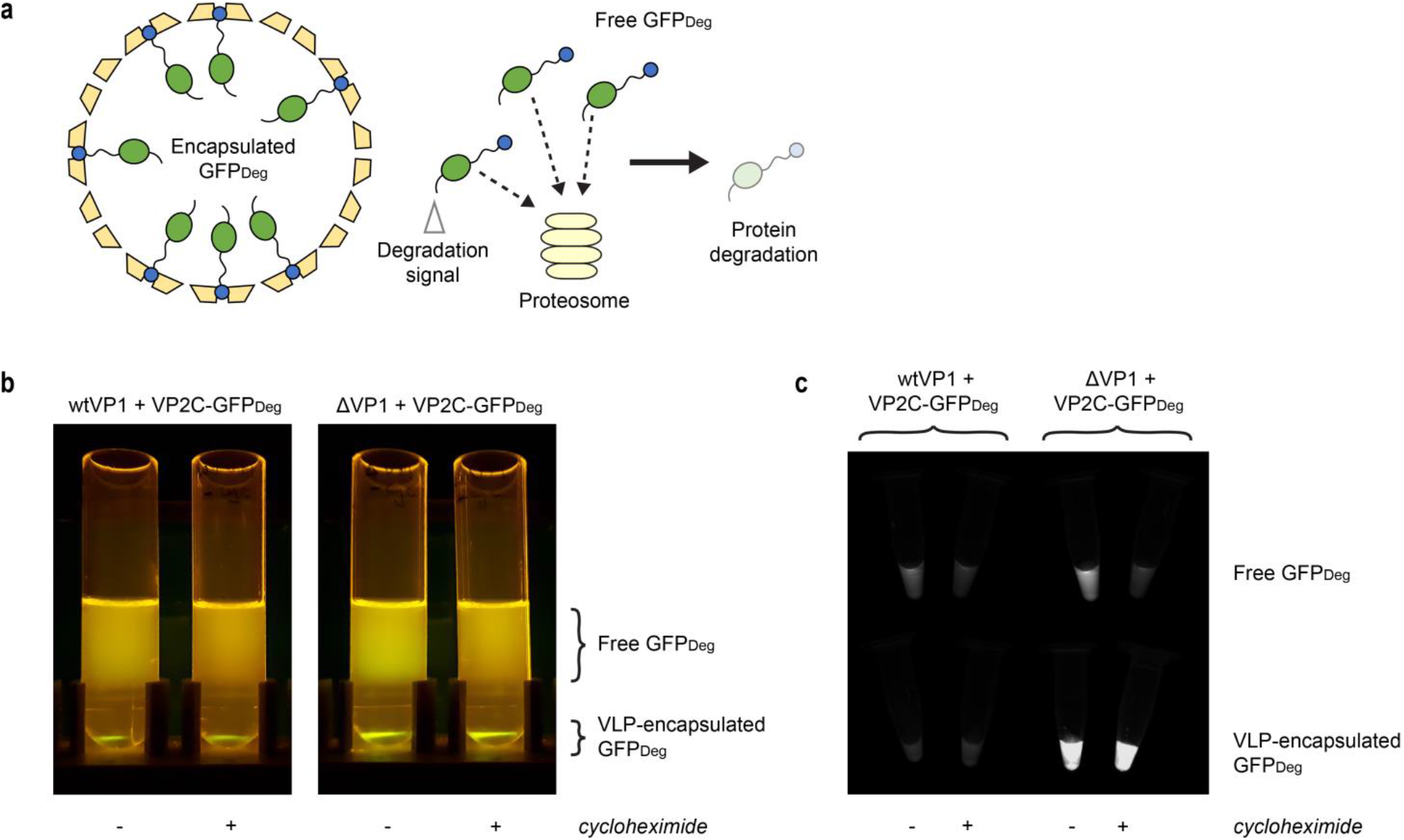
Destabilised GFP as a model protein for assessing *in vivo* compartmentalisation. (a) Tagging GFP with a degradation signal at the C-terminal (GFP_Deg_) targets it for proteasomal degradation, unless protected by VLP encapsulation. (b) Fluorescent lysates of VLP-expressing cells ultracentrifuged through an iodixanol cushion, with and without cycloheximide treatment. Substantial yeast autofluorescence can also be observed. (c) Samples collected from the ‘Free GFP_Deg_’ and ‘VLP-encapsulated GFP_Deg_’ layers from (b), viewed under blue light illumination and a 530 nm emission filter.

To investigate subcellular localisation *in situ*, cells were imaged by confocal laser scanning microscopy after 24 hours galactose induction and 3 hours cycloheximide treatment (Figure 4). The treatment duration was selected based on preliminary time-course experiments which showed that the residual GFP signal plateaus off ~2 hours after treatment (Figure S4). Each VP1 variant was coexpressed either with VP2C-GFP_Deg_, or GFP_Deg_ as a control without directed GFP encapsulation. ΔVP1 compartments appear to be distributed throughout the cell while wtVP1 led to the formation of small, localised foci either adjacent to or co-localised with the DNA stain (Figure 4). This is consistent with a previous study on yeast-expressed wtVP1, where clusters of assembled VLPs were found associated with tubulin fibres in the nucleus^11^. Importantly, it shows that the ΔVP1 variant redirects VLP compartment assembly and cargo loading to the cytosol, where there is access to a greater range of metabolites.

**Figure 4.**
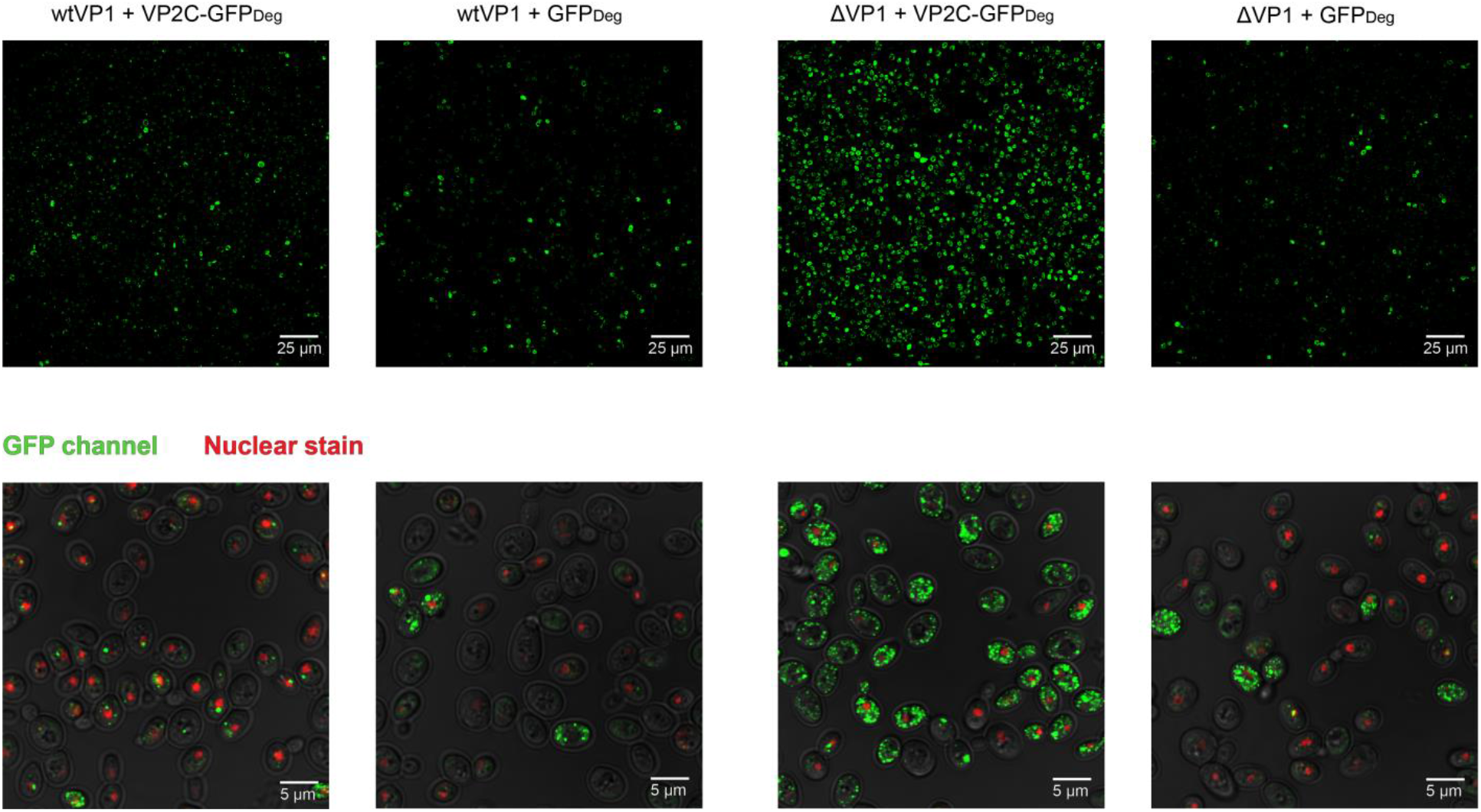
Visualising compartment localisation by confocal microscopy. Cells expressing GFP_Deg_ are imaged after 24 h galactose induction and 4 h cycloheximide treatment. The top panel shows a wide field of view while the bottom panel is 5X zoomed relative to the top. The top panel shows only the GFP channel (coloured green) while the bottom panel shows merged images of the GFP, nuclear stain (coloured red), and brightfield channels. The nuclear stain is Hoechst 34580, which is specific to dsDNA. The same imaging and processing parameters are used for all samples. To reduce autofluorescence, brightness and contrast for the GFP channel was adjusted until minimal signal is visible in the untransformed negative control strain (Figure S5).

The higher intensity of GFP fluorescence for ΔVP1 + VP2C-GFP_Deg_ indicates better cargo capture compared to wtVP1 + VP2C-GFP_Deg_ (Figure 4), consistent with ultracentrifugation observations (Figure 3b). This may be due to co-localisation of ΔVP1 with VP2C-GFP in the cytoplasm during assembly, which increases the probability of VP1-VP2C interactions. VP1 expression levels were similar for wtVP1 and ΔVP1, and were stable up to at least 72 hours post-induction (Figure S6). In contrast to a previous report showing that MPyV wtVP1 overexpression in yeast leads to temporary growth inhibition^11^ and despite being expressed with the strong GAL1 promoter, neither VP1 variant negatively impacted growth rates under the conditions tested (Figure S6). This is desirable because non-target physiological effects should be avoided both for examining basic biology and for applying synthetic biology tools in an industrial setting.

The confocal imaging indicated that there is a strong protective effect for encapsulated cargo. Flow cytometry was used to investigate this effect with increasing induction time by tracking the level of GFP_Deg_ before and after cycloheximide treatment (Figure 5). We only characterised the ΔVP1 variant because it exhibited preferred properties, namely cytoplasmic localisation and better cargo protection. The GFP_Deg_ encapsulation strain (ΔVP1 + VP2C-GFP_Deg_) was compared with controls lacking either VP1 or the VP2C anchor. At every time point, the signal after cycloheximide treatment (‘After cyc’) of the GFP encapsulation strain was significantly higher than that of the controls, indicating the stabilisation of a proportion of GFP_Deg_ from degradation. The majority of cargo proteins were unencapsulated in the early induction phase (< 24 hours), as indicated by the large difference in signal before and after cycloheximide treatment. As protein synthesis slowed down, the proportion of encapsulated GFP_Deg_ relative to total GFP_Deg_ increased. From 24 hours, all GFP_Deg_ appears to be encapsulated for ΔVP1 + VP2C-GFP_Deg_ while GFP_Deg_ depletion continued in the controls.

**Figure 5.**
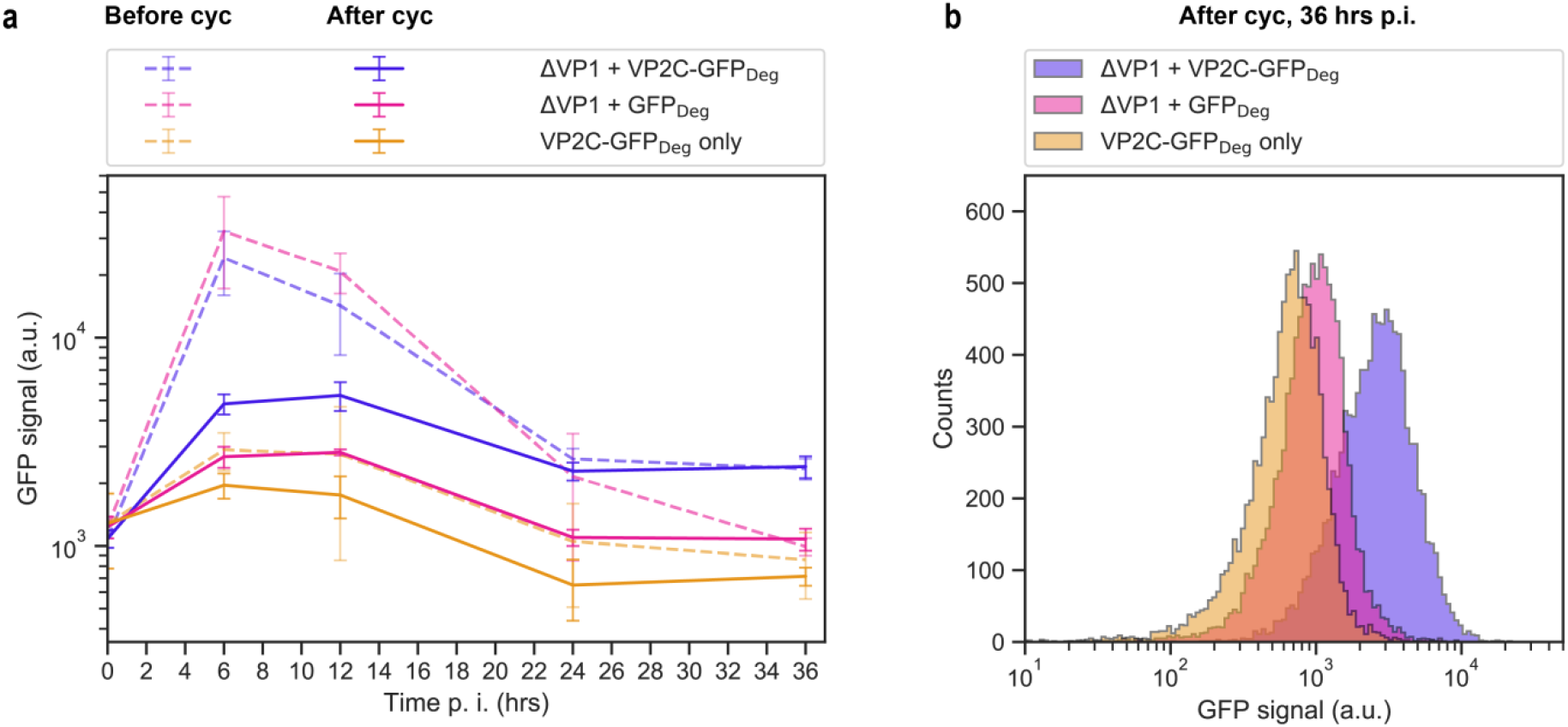
Investigating the protection of destabilised GFP by flow cytometry. (a) Flow cytometry of GFP_Deg_ strains before and after 3 h cycloheximide treatment (‘Before cyc’ and ‘After cyc’ respectively). The median was used to represent each sample of 10,000 cells. Values shown in the plot are the mean of 3 biological replicates, +/− 1 STD. (b) Sample raw flow cytometry histograms showing the population distribution for cycloheximide-treated cells at 36 h post-induction.

The difference in signal between strains expressing GFP_Deg_ and VP2C-GFP_Deg_ in the absence of VP1 (Figure 5, Figure S4) suggests that VP2C itself also destabilises the cargo protein prior to encapsulation. During prokaryotic expression, the N-terminal fusion renders GFP insoluble^14^; however, this is rescued by VP1 co-expression as VP2C-VP1 binding masks the hydrophobic motif present on VP2C^22^. Despite the reduced stability of VP2C-tagged GFP evident here in yeast, the effect of specific encapsulation led to a higher level of persistent GFP_Deg_ than the control lacking VP2C.

### Artificial compartmentalisation as a novel enzyme stabilisation strategy

After parameterising the system, we next examined if enzymes encapsulated *in vivo* by MPyV remain functional and can participate in a bioproduction pathway. At the same time, we sought to explore a novel *in vivo* use for self-assembling protein compartments in metabolic engineering as a general-purpose platform for stabilising enzymes. We identified the cytoplasmic enzyme myo-inositol oxygenase (MIOX) as a suitable target for encapsulation due to its apparent instability when expressed in heterologous hosts^39,40^. An artificial pathway has been described that only requires two enzymes to convert myo-inositol into D-glucaric acid, namely *Mus musculus* MIOX and *Pseudomonas syringae* uronate dehydrogenase (UDH)^39^. This production pathway has previously been expressed and shown to be functional in *S. cerevisiae*^41^. MIOX is the rate limiting enzyme of the pathway; subsequent engineering efforts for D-glucaric acid production have focused on improving its intracellular stability^42^ and expression level^40^. Furthermore, mouse MIOX is a small, monomeric protein^43^ and appears to be generally tolerant to fusions at the N- and C-termini^42,44,45^, which makes it an ideal candidate for exploring encapsulation within MPyV compartments via fusion to the self-sorting anchor, VP2C.

ΔVP1 and VP2C-MIOX were expressed using galactose-inducible promoters (P_GAL1_ and P_GAL10_) while the second enzyme in the pathway, UDH was expressed using the strong constitutive TEF1 promoter (Figure 6a, refer to Table 1 in Methods for strain details). A set of constructs were also generated with GFP fused at the N-terminus of MIOX as a reporter for flow cytometry and western blot. Altogether, five MIOX expression strategies were evaluated: ΔVP1 + VP2C-MIOX (VLP-forming), MIOX only (free control), ΔVP1 + VP2C-GFP-MIOX (VLP with GFP fusion), GFP-MIOX only (free control with GFP fusion), and VP2C-GFP-MIOX only (free control with VP2C anchor and GFP fusion). All expression cassettes were integrated as single copies in the yeast genome to ensure stable and uniform gene expression. As per previous studies^40,41^, myo-inositol was supplied directly in the culture medium and products were measured by sampling the culture medium (Figure 6b). Cultures were transferred from a glucose-containing medium to a galactose-containing medium upon flask inoculation, so the time of inoculation could be considered the point of ΔVP1 and MIOX induction.

**Table 1.** *S. cerevisiae* strains for D-glucaric acid production.

| Strain | Genotype | Reference |
| --- | --- | --- |
| CEN.PK2-1C | MATa ura3-52 leu2-3,112 trp1-289 his3Δ1 MAL2-8c SUC2 | Entian and Kötter (2007)<sup>46</sup> |
| UDH only | CEN.PK2-1C derivative;<br>leu2::T<sub>CYC1</sub>-UDH-P<sub>TEF1</sub>-P<sub>KILEU2</sub>-KILEU2 | This work |
| ΔVP1 + VP2C-MIOX | UDH only derivative;<br>ura3::KIURA3-T<sub>HIS5</sub>-(VP2C-MIOX)-P<sub>GAL10</sub>-P<sub>GAL1</sub>-ΔVP1 | This work |
| MIOX only | UDH only derivative;<br>ura3::KIURA3-T<sub>HIS5</sub>-MIOX-P<sub>GAL10</sub>-P<sub>GAL1</sub> | This work |
| ΔVP1 + VP2C-GFP-MIOX | UDH only derivative;<br>ura3::KIURA3-T<sub>HIS5</sub>-(VP2C-GFP-MIOX)-P<sub>GAL10</sub>-P<sub>GAL1</sub>-ΔVP1 | This work |
| GFP-MIOX only | UDH only derivative;<br>ura3::KIURA3-T<sub>HIS5</sub>-(GFP-MIOX)-P<sub>GAL10</sub>-P<sub>GAL1</sub> | This work |
| VP2C-GFP-MIOX only | UDH only derivative;<br>ura3::KIURA3-T<sub>HIS5</sub>-(VP2C-GFP-MIOX)-P<sub>GAL10</sub>-P<sub>GAL1</sub> | This work |

\* KIURA3 = URA3 from *Kluyveromyces lactis*, and KILEU2 = LEU2 from *Kluyveromyces lactis*.

**Figure 6.**
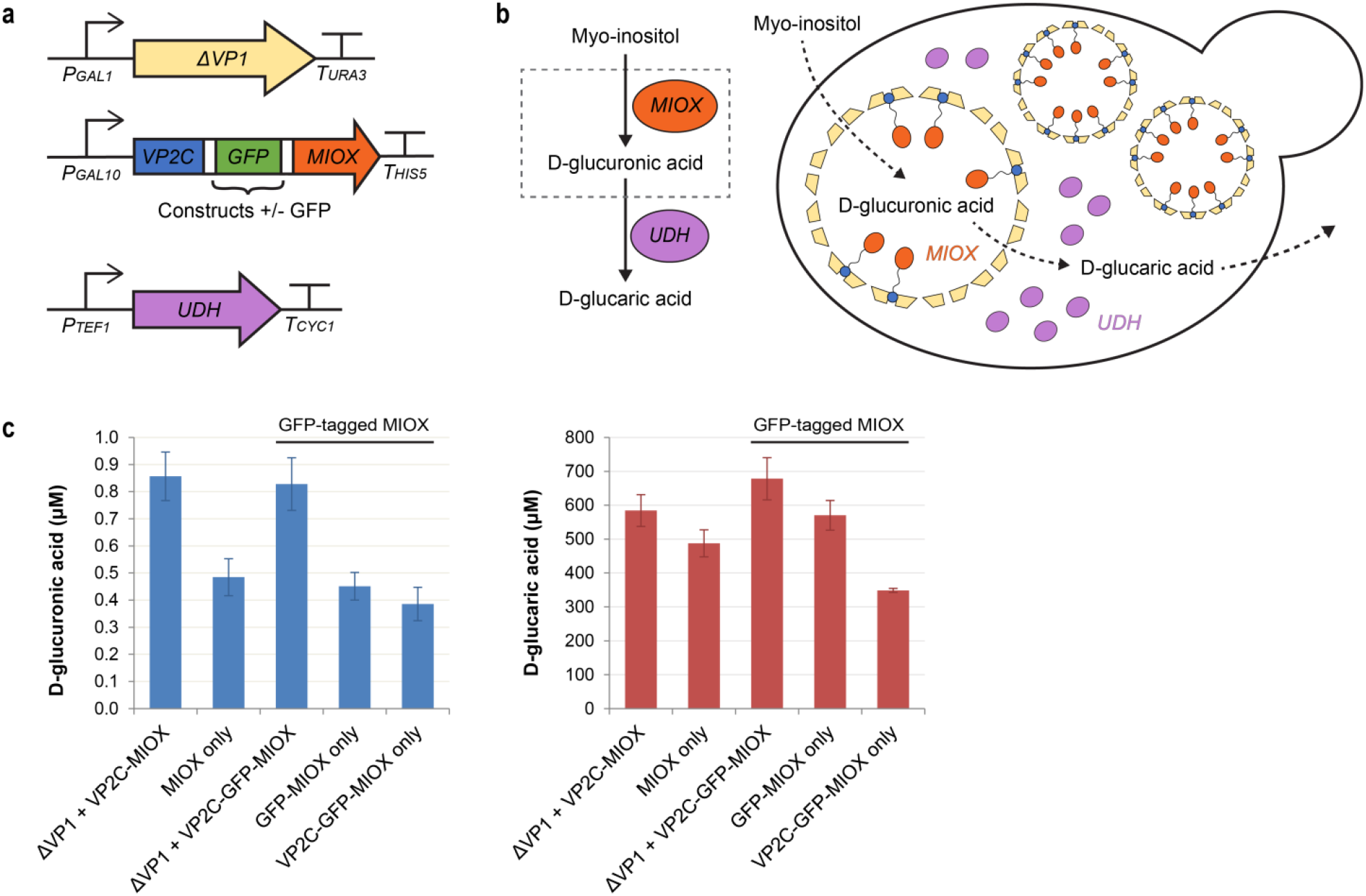
Compartmentalisation of myo-inositol oxygenase (MIOX) improved D-glucaric acid production. (a) Expression cassettes for MIOX encapsulation. Constructs were made with and without GFP as part of the cargo fusion protein. UDH is expressed using a strong constitutive promoter (P_TEF1_). All genes were chromosomally integrated as a single copy. (b) Reaction schematic of the heterologous D-glucaric acid pathway and an illustration showing proposed metabolite movements in the cell. The dashed grey box represents the compartment. Myo-inositol in the culture medium is taken up by yeast cells and diffuses into VLPs. Encapsulated MIOX converts myo-inositol into D-glucuronic acid, which then diffuses out into the cytoplasm where it is further converted into D-glucaric acid by uronate dehydrogenase (UDH). D-glucaric acid is then released into the culture medium. (c) Final titres of D-glucuronic acid and D-glucaric acid in the culture medium after 72 h fermentation. All data points are the means of 3 biological replicates; error bars are +/− 1 STD.

The D-glucuronic acid (intermediate) and D-glucaric acid (end product) titres in the culture medium were quantified by gas chromatography coupled to mass spectrometry (GC-MS) at 72 h (Figure 6c). The much lower concentration of D-glucuronic acid detected compared to D-glucaric acid (~3 orders of magnitude difference) suggests that UDH activity is not limited in any of the strains, and indicates that D-glucuronic acid produced by MIOX could readily escape VLPs. Coexpression of ΔVP1 with VP2C-GFP-MIOX almost doubled the final D-glucaric titre (*p* < 0.001) compared to VP2C-GFP-MIOX alone. VLP-forming strains (with or without GFP fusion) produced ~20% more D-glucaric acid than their corresponding free MIOX controls without the destabilising VP2C anchor; the increases were however not statistically significant (*p* = 0.053 for MIOX pair and *p* = 0.070 for GFP-MIOX pair, two-tailed Student’s *t*-test) for the size of the dataset used. Interestingly, N-terminal fusion of GFP to free MIOX also increased D-glucaric acid production – presumably by improving protein stability. This is in contrast to a previous *E. coli* study^42^, where fusion of MBP to the N-terminus of MIOX caused loss of *in vivo* enzyme activity.

The final cell densities of the two VLP-forming strains were >40% higher compared to the free MIOX controls (Figure 7a), pointing to the mitigation of some form of cellular toxicity by compartmentalisation. This is an intriguing finding as neither myo-inositol nor D-glucaric acid were expected to be toxic at these concentrations: a previous yeast study found that extracellular concentrations of D-glucaric acid up to 5 g/L (23.8 mM) did not negatively affect strain growth or productivity^41^; similarly, they did not observe any difference in growth rates with or without supplementation of myo-inositol at the same concentration used here (60 mM)^41^. In contrast, the growth profiles of strains with encapsulated MIOX were similar to that of a non-MIOX-expressing control strain (Figure S7), suggesting that overexpression of the MIOX protein may be inherently toxic to yeast. This is similar to a recent yeast study where targeting of norcoclaurine synthase into peroxisomes was found to alleviate cellular toxicity associated with the enzyme^47^.

**Figure 7.**
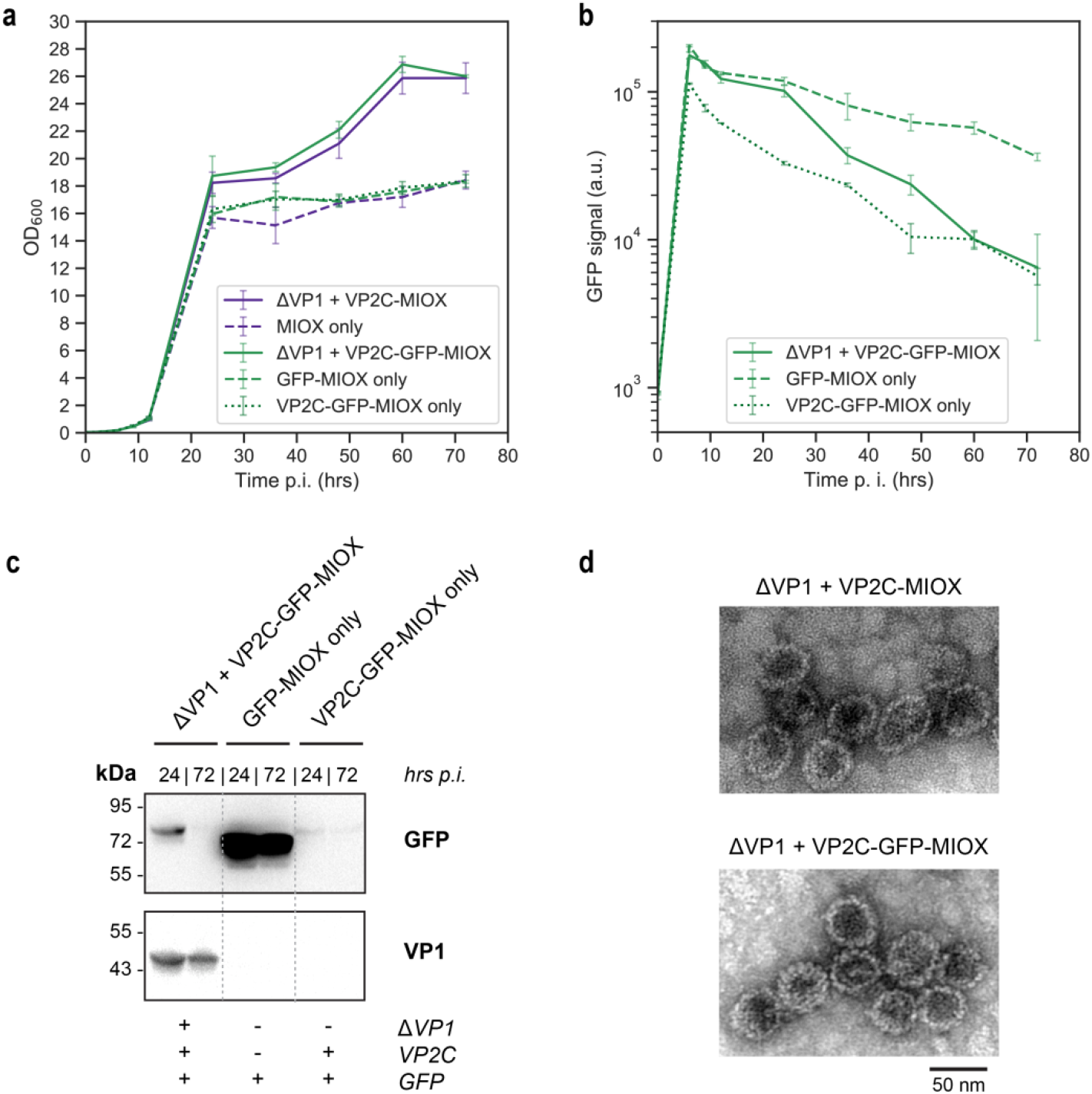
Growth profiles and protein expression levels of MIOX-expressing strains. (a) Cell density (OD_600_) against time post-induction. (b) GFP fluorescence was tracked by flow cytometry as a proxy for MIOX levels in the three GFP-tagged MIOX constructs. (c) Anti-GFP and anti-VP1 western blots of cell lysates at 24 h and 72 h post-induction. The same amount of cells was loaded per lane, based on the OD_600_ reading. Bands on the anti-GFP blot match the expected size of each corresponding MIOX fusion protein. (d) MIOX compartments isolated by iodixanol cushion ultracentrifugation, negatively stained and viewed under TEM. All data points in (a) and (b) are the means of 3 biological replicates; error bars are +/− 1 STD.

GFP signal detected by flow cytometry (Figure 7b) and western blot (Figure 7c) was used as a proxy for intracellular MIOX levels in the three GFP-tagged strains. The GFP-MIOX only strain had the most protein, followed by the ΔVP1 + VP2C-GFP-MIOX and VP2C-GFP-MIOX only strains. Sorting into compartments clearly increased the stability of VP2C-GFP-MIOX; however, the destabilisation effect of the VP2C anchor on cargo proteins was apparent – as observed earlier with GFP_Deg_ strains (Figure 5). Given the huge difference in amount of GFP-MIOX in the GFP-MIOX only and ΔVP1 + VP2C-GFP-MIOX, the increase in D-glucaric acid titre for the VLP-forming strain (Figure 6c) is quite remarkable. This result indicates a very strong stabilising effect of encapsulation on MIOX activity, which may be similar to the stabilising effect of encapsulation within protein cages observed for a number of enzymes^48,49^. Despite protection by compartmentalisation, VP2C-GFP-MIOX levels decreased after 24 h in the VLP-forming strains. This may be due to the proportion of cargo (i.e. VP2C-GFP-MIOX) captured by VLP assembly being relatively small compared to the total expressed cargo, as also observed in the GFP_Deg_ experiments (Figure 5a). Since active cell growth in the VLP strains continued beyond 24 h (after galactose would have been fully consumed), switching-off of galactose-inducible promoters may have additionally ‘diluted’ MIOX and ΔVP1 levels in the daughter cells to a greater degree compared to the other strains. VLP formation in ΔVP1-expressing constructs was verified by TEM (Figure 7d) and VLPs isolated from ΔVP1 + VP2C-GFP-MIOX were fluorescent green (data not shown), confirming the presence of cargo protein. Overall, the results show that the positive effect of MIOX encapsulation on growth and biomass accumulation was able to compensate for reduced enzyme levels in terms of D-glucaric acid production.

## CONCLUSIONS

We have established the engineered MPyV VLP system as a simple and orthogonal platform for protein compartmentalisation in yeast. Implementation of a cytoplasm-localised shell variant (ΔVP1) allowed specific and efficient compartmentalisation of cytoplasmic cargo proteins. We then explored the compartmentalisation of a naturally unstable metabolic enzyme, MIOX, for the bioproduction of D-glucaric acid. In contrast to previous *in vivo* studies which used protein compartments as a scaffold for co-localising multiple enzymes in a reaction cascade^4,5^, we wanted to investigate if a pathway can be improved simply by encapsulating a single, rate-limiting enzyme. Strains with encapsulated MIOX successfully produced D-glucaric acid at higher titres than free MIOX. This is the first demonstration in yeast of a synthetic biocatalytic compartment that can participate in a metabolic pathway and shows that metabolites can diffuse through the MPyV shell. Moreover, an increased target product titre was achieved despite dramatically lower levels of expressed protein. Compartment-forming strains grew to higher cell densities than the free controls, suggesting a protective effect against cellular toxicity. This work also provides proof-of-concept of using an orthogonal self-assembling protein compartment for protecting metabolic enzymes from *in vivo* degradation.

To extend the work on D-glucaric acid production, it may be useful to investigate if deletion of the major inositol pathway regulator OPI1 could improve titres by increasing intracellular myo-inositol^40,41^. Alternatively, co-encapsulation of MIOX with the downstream enzyme, UDH, may improve accessibility to the intermediate D-glucuronic acid. Protein co-encapsulation with the MPyV compartment in yeast is the subject of ongoing work. Another key direction for artificial *in vivo* metabolons like MPyV VLPs will be to maximise the proportion of encapsulated cargo proteins. Our results show that encapsulation in self-assembling compartments is a promising strategy for isolating individual nodes in a reaction pathway and shielding proteins from specific interactions with host factors. Ultimately, we envision this self-assembling compartment as a versatile ‘plug-and-play’ tool for studying and harnessing *in vivo* catalysis.

## METHODS

### Molecular cloning and strain generation

All cloning was performed using the isothermal assembly method, using NEBuilder HiFi DNA Assembly Master Mix (NEB #E2621). ΔVP1 was codon-optimised for yeast and synthesised by GenScript. All other synthetic genes were manually codon-optimised for *S. cerevisiae* and synthesised as dsDNA fragments by Integrated DNA Technologies. First, the ΔVP1 empty plasmid was constructed by replacing the GFP sequence in pILGFPB5A^50^ (YIp with a *K. lactis* URA3 marker) with PCR-amplified ΔVP1. Then, the ΔVP1 + VP2C-GFP construct was generated by inserting PCR-amplified VP2C and GFP fragments between the XbaI and EcoRI sites of the ΔVP1-only plasmid. All other constructs (except UDH cassette) were made by replacing either ΔVP1 or VP2C-GFP of these two plasmids by restriction digest, gel purification, and isothermal assembly. PCR primer and synthetic gene sequences are listed in Table S1 and Table S2 respectively. GFP was swapped for different cargo proteins (GFP_Deg_, MIOX, GFP-MIOX) by double digesting with BamHI and BglII. For control constructs without VP1, VP1 was excised by NotI and NheI (NEB) digestion and patched with a single-stranded DNA oligonucleotide. After incubating at 50 °C for 1 hour, each assembly reaction mix was directly transformed into chemically competent *E. coli* DH5α by heat-shock. Plasmids purified from individual colonies were verified for the correct insert by Sanger sequencing (Australian Genome Research Facility). The PTEF1-UDH-TCYC1 expression cassette was generated over multiple assembly steps, starting with the pUG73^51^ backbone vector which contains a *Kluyveromyces lactis* LEU2 marker.

For all constructs, the plasmids were first digested with SwaI (NEB #R0604) and transformed using the LiAc/SS carrier DNA/PEG method^52^, leading to stable single-copy integration into the yeast genome. The base strain is CEN.PK2-1C (MATa, his3D1, leu2-3_112, ura3-52, trp1-289, MAL2-8c, SUC2)^46^ (Euroscarf). For MIOX and UDH coexpression (Table 1), the base strain was first transformed with the P_TEF1_-UDH-T_CYC1_ cassette (contains a leucine auxotrophic selection marker, LEU2). A single transformant was then used for the second transformation with MIOX expression cassettes. Yeast transformants were verified by colony PCR using the same primers used for cloning. Refer to Table S3 for strain, plasmid, and protein part details. For every construct, at least three colonies recovered from yeast transformation were selected and maintained as biological replicates. Strains were grown overnight in YPD (2% w/v Bacto Peptone, 1% w/v Bacto yeast extract, 2% w/v glucose) and stored as 20% v/v glycerol stocks at −80 °C.

### VLP expression and purification

All incubations were performed at 30 °C, 200 rpm shaking (Infors HT Multitron incubator). Glycerol stocks were recovered on uracil drop-out agar plates and pre-cultured overnight in YPD. YPD cultures were diluted into YPGD (2% w/v Bacto Peptone, 1% w/v Bacto yeast extract, 2% w/v galactose, 0.5% w/v glucose) at OD_600_ = 0.2 and grown for 24 hours. For VLP purification, we routinely grew 200-300 ml cultures in 500 ml unbaffled shake flasks. Cells were collected by centrifugation and stored at −20 °C until required.

Thawed cell pellets were resuspended in lysis buffer (20 mM MOPS, 150 mM NaCl, 1 mM CaCl_2_, 0.01% Triton X-100, pH 7.8) and lysed in 3 passes at >22,000 psi with a high-pressure homogeniser (Avestin Emulsiflex C5). The PEG-NaCl method^53^ was used as a concentration and initial purification step. Briefly, NaCl and PEG 6000 were added to a final concentration of 0.5 M and 8% w/v respectively (from a 5X PEG-NaCl stock). After storing overnight at 4 °C, the precipitate was collected by centrifugation and resuspended in 2 ml of Buffer A (20 mM MOPS, 150 mM NaCl, 1 mM CaCl_2_, pH 7.8). 1-2 ml PEG-concentrated sample or clarified lysate was layered onto a 1 ml cushion of 30% iodixanol (OptiPrep) in Buffer A. Ultracentrifugation was run for 3 hours at 100,000 g, 8 °C (Beckman Coulter Optima MAX-XP, TLA-100.3 fixed-angle rotor). 100-200 μl of VLP sample was collected from the ‘dense’ fraction at the bottom of each tube. Note: iodixanol absorbs strongly in the UV range and interferes with TEM negative staining – buffer exchange or sample dilution is advisable before further analysis.

The ultracentrifugation step isolates VLPs along with two high-MW yeast contaminants (Figure S8). An SEC step was used to polish samples and remove iodixanol. Samples were topped up to 1 ml with Buffer A and loaded onto a HiPrep 16/60 Sephacryl S-500 HR column (GE Healthcare). Buffer A was run at 1 ml/min and the flow-through was collected in 5 ml fractions. MPyV VLPs elute as a broad peak around 50-70 ml (Figure S8c). VLP fractions were pooled and concentrated using 100 kDa MW cut-off centrifugal filters (Amicon Ultra 4 ml, Merck). Protein concentrations were measured using the linearised Bradford method^54^ with Pierce Coomassie Protein Assay Kit (Thermo Scientific).

### Gel electrophoresis

Three μg purified VLPs was loaded per lane. SDS-PAGE was run on Any kD Mini-PROTEAN TGX gels (Bio-Rad) in Tris-glycine SDS running buffer (25 mM Tris, 192 mM glycine, 0.1% w/v SDS, pH 8.3) at 150 V, 55 min and stained with GelCode Blue Safe Protein Stain (Thermo Scientific #24594). The protein MW marker used was Blue Prestained Protein Standard, Broad Range (11-250 kDa) (NEB #P7718). Native gel electrophoresis of intact VLPs was run on a 1% w/v agarose mini gel (7×7 cm) in TA buffer (40 mM Tris-base, 20 mM acetic acid) at 90 V for 60 min. VLP samples were suspended in buffer with bromophenol blue and 10% v/v glycerol (final concentration) to aid loading. EDTA was avoided in buffers as polyomavirus VLPs are known to be stabilised by interactions with calcium ions^30,55^. Nucleic acid staining was performed by soaking the gel in 1X GelRed (Biotium #41003) in TA buffer for 1 hour at room temperature, with gentle shaking. 0.5 μg DNA ladder (Thermo Scientific #SM0311) was loaded as a positive control. For visualising proteins, gels were stained with GelCode Blue Safe Protein Stain (Thermo Scientific #24594) overnight. Images were captured with a ChemiDoc MP Imaging System (Bio-Rad). Imaging settings: ‘Fluorescein’ preset (blue epi excitation, 530/30 nm filter) for GFP fluorescence, ‘Ethidium bromide’ preset (UV excitation, 605/50 nm filter) for stained nucleic acid, ‘Coomassie blue’ preset for protein.

### Transmission electron microscopy (TEM)

VLP samples were diluted in phosphate buffered saline (PBS) or Buffer A to ~0.1 mg/ml and settled on formvar/carbon coated copper mesh grids (ProSciTech #GSCU200C) for 1-2 min. Grids were briefly rinsed in a drop of distilled water (excess removed with filter paper) and stained with 1% w/v aqueous uranyl acetate for 1 min. Uranyl acetate was then blotted off with filter paper and the grids air dried for a few minutes before storing. Grids were imaged with a Hitachi HT7700 transmission electron microscope at 80 kV (High Contrast mode).

### Nanoparticle tracking analysis (NTA)

VLP samples were diluted ~100 ng/ml in Buffer A and analysed with a NanoSight NS300 (Malvern Panalytical) equipped with a 405 nm laser and temperature control. Syringe pump speed during capture was set at 100 and 3 x 60s videos were recorded for each sample. Optimal particle concentration was 50-100 particles/frame; if required, samples were diluted further and re-analysed until the captured data falls within the acceptable range. Imaging settings: Camera Level = 15, Temperature = 25.0 °C, Viscosity = 1.0 cP, Detect Threshold = 5. Raw particle data was exported for further analysis.

### Analytical size-exclusion chromatography (SEC)

Separations were performed with a Shimadzu Prominence XR HPLC with a Nexera Bio Kit connected to MALS (Wyatt Dawn 8) and UV-Vis absorbance (Shimadzu SPD-M20A photodiode array) detectors. Purified VLP samples were diluted to 0.1-0.2 mg/ml in PBS and filtered with 0.22 μm cellulose acetate spin filters (Sigma-Aldrich #CLS8161). 25 μl of each sample was injected through a Bio SEC-5 2000 Å HPLC column (Agilent) with a Bio SEC-5 2000 Å guard (Agilent). The mobile phase was PBS, with a constant flow rate of 1 ml/min for a 20 min run. The laser wavelength for MALS was 659 nm. The molar mass was fitted based on a Zimm light scattering model using Wyatt ASTRA 7 software.

### Ultracentrifugation analysis

YPD overnight pre-cultures were diluted into 50 ml YPG at OD600 = 0.4 and grown for 6 hours at 30 °C, 200 rpm shaking. Cycloheximide was added to 100 μg/ul and cultures were returned to the incubator for a further 3 hours to allow sufficient degradation of unencapsulated cargo protein. Cells were collected by centrifugation. The same amount of cells for each sample was transferred to 2 ml screw-capped tube, adjusting based on the OD_600_ value. The samples were resuspended to 1 ml total volume in lysis buffer and vortexed with ~0.5 g 0.5 mM glass beads on a tabletop vortex mixer with a microtube rack. Six cycles of 1 min vortex + 1 min on ice were performed. Debris was removed by centrifugation at 12,000 g for 5 min. The clarified lysate was layered onto a 1 ml cushion of 30% iodixanol (OptiPrep) in Buffer A in clear ultracentrifuge tubes (3.5 ml thickwall polycarbonate tubes, Beckman Coulter #349622). Ultracentrifugation was run for 3 hours at 100,000 g, 8 °C (Beckman Coulter Optima MAX-XP, TLA-100.3 fixed-angle rotor). Ultracentrifuge tubes were photographed through an orange filter, backlit with a blue light transilluminator (Safe Imager 2.0, Invitrogen). 200 μl samples were collected from the top ‘free GFP_Deg_’ and dense ‘VLP-encapsulated GFP_Deg_’ layers into clear 1.5 ml microcentrifuge tubes. All tubes were imaged simultaneously with a ChemiDoc MP Imaging System (Bio-Rad) using the ‘Fluorescein’ preset (blue epi excitation, 530/30 nm filter).

### Flow cytometry

YPD overnight pre-cultures were diluted 1:100 into 3 ml YPG (2% w/v Bacto Peptone, 1% w/v Bacto yeast extract, 2% w/v galactose) in 24-well culture plates and grown at 30 °C, 200 rpm shaking. Flow cytometry was performed on live cells immediately after sampling using an Accuri C6 Flow Cytometer (BD Biosciences). At every time point, 500 μl culture was also transferred into a separate well containing cycloheximide (final concentration 100 μg/ml). Cultures with cycloheximide were returned to the incubator for a further 3 hours before flow cytometry. GFP signal was measured using 488 nm laser excitation and a 533/30 nm BP emission filter. 10,000 cells were sampled for each reading (trigger threshold FSC-H > 250,000). Each biological replicate is a separate colony recovered during yeast transformation.

### Confocal microscopy

YPD overnight pre-cultures were diluted into YPG at OD_600_ = 0.2 and grown for 24 hours at 30 °C, 200 rpm shaking. Cycloheximide was added to a final concentration of 100 μg/ul and cultures were returned to the incubator for a further 3 hours to allow sufficient degradation of unencapsulated cargo protein. Cells were harvested by gentle centrifugation, washed once with PBS and fixed with 4% w/v methanol-free formaldehyde (Thermo Scientific #28906) for 20 min at room temperature. Nuclear staining was performed by incubating cells with 10 μg/ml Hoechst 34580 (Invitrogen #H21486) for 30 min at room temperature. Cells were immobilised on glass-bottom dishes (Cellvis #D35C4-20-1.5-N) pre-coated with 0.1 mg/ml concanavalin-A (Sigma-Aldrich #C2010). Images were taken with an Olympus FV3000 confocal laser scanning microscope with a 60x silicone oil immersion objective (1.3 NA) using the ‘EGFP’ and ‘Hoechst 33342’ filter presets. Cells were focused using the Hoechst channel to minimise GFP photobleaching. Image brightness and contrast were adjusted using ImageJ and were kept consistent across the whole sample set. The untransformed base strain (CEN.PK2-1C) was used as a control for cell autofluorescence (see Figure S6).

### Western blot

Amount loaded per lane was normalised by OD_600_ readings (equivalent to 10 μl of culture at OD_600_ = 20). Samples were run on Any kD Mini-PROTEAN TGX gels (Bio-Rad) in Tris-glycine-SDS buffer at 150 V, 55 min and transferred onto nitrocellulose membranes (Amersham Protran 0.45 μm, GE Healthcare) by wet transfer at 75 V, 60 min. The transfer buffer was 1X SDS-PAGE buffer + 20% v/v methanol. Even protein transfer was verified by staining the membrane with 0.1% w/v Ponceau S in 5% v/v acetic acid. Membranes were blocked with 5% w/v skim milk in PBS + 0.05% v/v Tween 20 for >1h at RT and incubated with primary antibodies overnight at 4 °C. Membranes were washed briefly with blocking buffer and incubated with secondary antibodies for 90 min at RT. The antibodies and dilutions used were as follows: rabbit anti-VP1 antiserum 1:2000, mouse anti-GFP monoclonal IgG (Cell Signaling Technology #2955) 1:2000; anti-rabbit IgG-HRP (Cell Signaling Technology #7074) 1:2000, anti-mouse IgG-HRP (Cell Signaling Technology #7076) 1:2000. Rabbit anti-VP1 antiserum was produced by Walter and Eliza Hall Institute Antibody Services using wtVP1 expressed in *E. coli* and assembled into VLPs *in vitro*^18^. Blots were visualised with Clarity Western ECL Substrate (Bio-Rad #1705060) and imaged with a ChemiDoc MP Imaging System (Bio-Rad).

### Fermentation for D-glucaric acid production

Glycerol stocks were recovered on uracil drop-out plates and pre-cultured overnight in YPD. Cultures were inoculated to OD_600_ = 0.05 in 20 ml YPG + 60 mM myo-inositol, in 50 ml unbaffled shake flasks. ‘Time post-induction’ was counted from the time of inoculation. Cultures were grown at 30 °C with 200 rpm shaking. At every time point, cultures were sampled for OD_600_ and flow cytometry measurements. Culture samples for western blot and GC-MS were stored at −20 °C until further use.

### Metabolite analysis by GC-MS

Frozen cultures were thawed completely and centrifuged to pellet cells and debris. For each sample, 40-120 μl of culture supernatant was dried in 1.5 ml microcentrifuge tubes with a rotational vacuum concentrator (Concentrator Plus, Eppendorf) and derivatised in 20 μl methoxyamine chloride (30 mg/ml in pyridine) with continuous shaking for 2 h at 37 °C. 20 μl of *N,O*-bis(trimethylsilyl)trifluoroacetamide (BSTFA) containing 1% trimethylchlorosilane (TMCS) was then added and the samples further incubated for 30 min at 37 °C, with continuous shaking. Analytical standards (dissolved in distilled water) were derivatised the same way as the samples. The standards used were D-saccharic acid (D-glucaric acid) potassium salt >98% (Sigma-Aldrich #S4140) and D-glucuronic acid >98% (Sigma-Aldrich #G5269).

1 μl of the derivatised sample was injected into a PTV injector in splitless mode with an injection temperature of 250 °C. Separations were carried out in an Agilent J&W DB-5ms column (95% polydimethylsiloxane, 30 m x 0.25 mm x 1 μm) with a Shimadzu GC/MS-TQ8050 system. The GC system was equipped with an electron impact (EI) ionisation source and operated in multiple reaction monitoring (MRM) mode for the detection of compounds. The temperature of the oven was initially set at 120 °C and increased to 230 °C at a rate of 18 °C/min. The temperature was then increased at 8 °C/min until 265 °C and finally to 300 °C at 18°C/min. The total run time was 12.43 min. Helium was used as a carrier gas, with a constant flow rate of 1.0 ml/min. The ion source temperature was set at 230 °C and interface temperature was 280 °C. The specific mass spectrometric parameters were tailored for each compound individually in order to monitor the fragmentation ions for each analyte. Two fragment ions from the EI ion source were nominated and then further fragmented in the collision cell. The three most abundant transients were monitored. The first transient was used as the quantifier and the other two as qualifiers (see Table 2).

**Table 2.** Optimised MRM transitions and retention times for selected compounds analysed by GC-MS/MS.

| Compound | Retention time (min) | Transient 1 (m/z) | Collision Energy (V) | Transient 2 (m/z) | Collision Energy (V) | Transient 3 (m/z) | Collision Energy (V) |
| --- | --- | --- | --- | --- | --- | --- | --- |
| D-glucuronic acid | 10.348 | 333.0>73.1 | 29 | 160.0>73.1 | 17 | 333.0>143.1 | 13 |
| D-glucaric acid | 11.165 | 333.0>73.1 | 25 | 333.0>143.1 | 13 | 292.0>73.1 | 25 |

### Data analysis

Data analysis and plotting for NTA, SEC-MALS, OD_600_, and flow cytometry was performed with Python 3. Graphs were generated using the Matplotlib package. For NTA, raw particle data from each 60s capture was converted into a histogram (0.01 nm binwidth) and converted into a probability distribution by Gaussian kernel density estimation (KDE) using the scipy.stats package. The mean and standard deviation at every point of the 3 KDE curves in each sample were then calculated to generate the final distribution. For flow cytometry, raw flow cytometry data (.fcs files) was analysed with the FlowCal package. The median of each population of 10,000 cells (calculated using FlowCal) is used to represent each biological replicate. The mean and standard deviation of the 3 median values were then calculated to generate the final data points.

## ASSOCIATED CONTENT

Supporting information file (PDF).

Plasmids used in this work are available from Addgene (accession numbers to be provided upon manuscript acceptance).

## ACKNOWLEDGEMENTS

Li Chen Cheah is supported by a Commonwealth Research Training Program scholarship. Additional funding for this project was provided through a Commonwealth Scientific and Industrial Research Organisation (CSIRO) Synbio PhD Top-up Scholarship. Frank Sainsbury acknowledges support from CSIRO in the form of a Synthetic Biology Future Science Platform Fellowship. This work was supported in part by a Research Grant from the Human Frontier Science Program (Ref.-No: RGP0012/2018). The authors thank Donna McNeale (Griffith University) for preparing purified wtVP1 for raising VP1 antiserum and Mitchell O’Sullivan (Queensland University of Technology) for assistance with writing Python scripts for data analysis. The authors acknowledge the Translational Research Institute Australia (TRI) for providing the Microscopy Core Facility that enabled this research and thank Adler Ju for confocal microscopy technical assistance. The authors acknowledge the facilities, and the scientific and technical assistance, of the Microscopy Australia Facility at the Centre for Microscopy and Microanalysis (CMM), The University of Queensland. Aspects of this research have been facilitated by access to Metabolomics Australia and the Australian Proteome Analysis Facility, supported under the Australian Government’s National Collaborative Research Infrastructure Strategy (NCRIS).

