## Supplementary information for "An artificial self-assembling nanocompartment for organising metabolic pathways in yeast"

**List of Supplementary Figures and Tables:**

Figure S1. Yeast-expressed VP2C-cargo undergoes spontaneous truncation in vivo. 2

Figure S2. MPyV wtVP1 binds to and co-purifies with a short VP2C truncation variant when coexpressed in E. coli. 3

Figure S3. The VP2C anchor is required to direct encapsulation of the cargo protein. 4

Figure S4. Flow cytometry to optimise the duration of cycloheximide treatment. 5

Figure S5. Confocal microscopy images of strains expressing VP2C-GFP$_{Deg}$ alone and a blank untransformed strain (negative control). 6

Figure S6. Examining the effect of wtVP1 and ΔVP1 expression of cell growth and protein expression. 7

Figure S7. Compartmentalisation rescues growth defects from MIOX overexpression. 8

Figure S8. SEC polishing step for VLP purification. 19

Table S1. PCR primer sequences. 9

Table S2. Synthetic gene sequences. 13

Table S3. Yeast strain, plasmid, and protein details. 16

Figure S1. Yeast-expressed VP2C-cargo undergoes spontaneous truncation *in vivo*.

(a) Anti-VP1 and anti-GFP western blots on purified VLPs to verify band identity. Total protein was stained using Ponceau S.

(b) Sites of VP2C truncation, as determined by N-terminal sequencing (Edman degradation). The N-terminal residues of each of the two SDS-PAGE cargo bands are underlined. The larger SDS-PAGE product ('upper band') is the full length cargo (except for removal of the first methionine), and the smaller product ('lower band') is a degradation product cleaved between Tyr-23 and Glu-24 of VP2C.

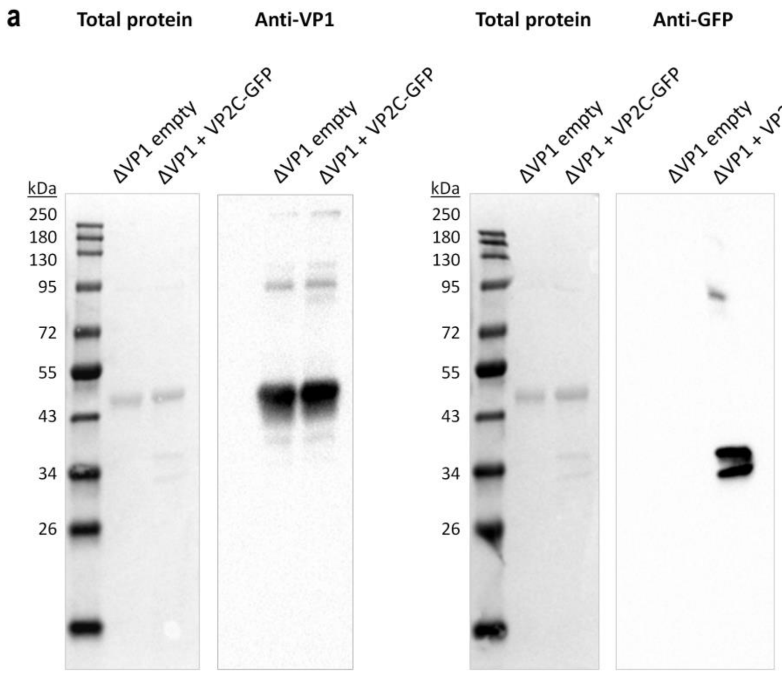


b VP2C-linker-GFP sequence:

MGNGGPTPAAHIQDESGEVIKFYQAQVVSHQRVTPDWMLPLILGLYGDITPG
GGGSGGGGSGGGGSMGSSKGEELFTGVVPILVELDGDVNGHKFSVSGEGEGD
ATYGKLTLKFICTTGKLPVPWPTLVTTFGYGVQCFARYPDHMKQHDFFKSAM
PEGYVQERTIFFKDDGNYKTRAEVKFEGDTLVNRIELKGIDFKEDGNILGHK
LEYNYNSHNVYIMADKQKNGIKVNFKIRHNIEDGSVQLADHYQQNTPIGDGP
VLLPDNHYLSTQSALSKDPNEKRDHMVLLEFVTAAGITHGMDELYKRS

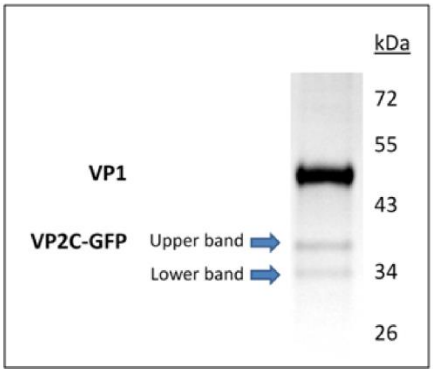

Figure S2. MPyV wtVP1 binds to and co-purifies with a short VP2C truncation variant when coexpressed in *E. coli*.

(a) GFP was used as the model cargo protein and fused translationally to VP2C at the N-terminal and a 6His tag at the C-terminal. The 23-residue variant of VP2C starts at the serine 5 residues after the Glu-24 underlined in Figure S1b. VP2C-GFP-6His and wtVP1 were coexpressed using a pETDuet vector with dual T7 promoters, as described previously[1].

(b) Purification of His-tagged VP2C-GFP from cell lysate by immobilised metal affinity chromatography (IMAC). Note that for *E. coli* expression, MPyV VP1 forms capsomeres but does not further self-assemble into VLPs *in vivo*[1], leaving the His tag exposed and available for binding to the IMAC column.

(c) Analysis of IMAC elution fractions from (b) by SDS-PAGE and anti-VP1 dot blot shows that VP1 co-purifies with the 23-residue VP2C variant.

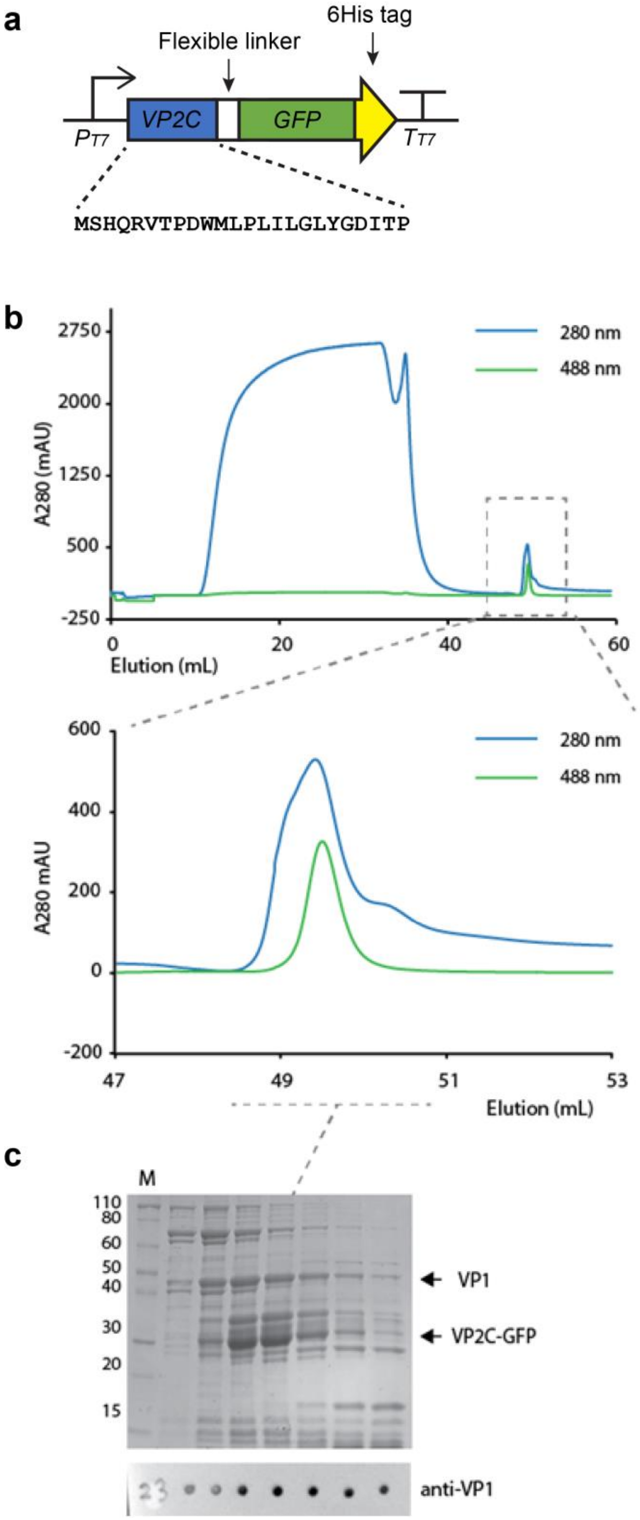

Figure S3. The VP2C anchor is required to direct encapsulation of the cargo protein.

Coexpression of VP1 with $GFP_{Deg}$ without VP2C led to absence of fluorescence in the dense particle fraction after ultracentrifugation of cell lysate through a 30% iodixanol cushion. The two tubes for each construct were loaded with different cell densities (based on $OD_{600}$) prior to ultracentrifugation. Tubes and samples were illuminated with blue light and viewed with a 530 nm filter.

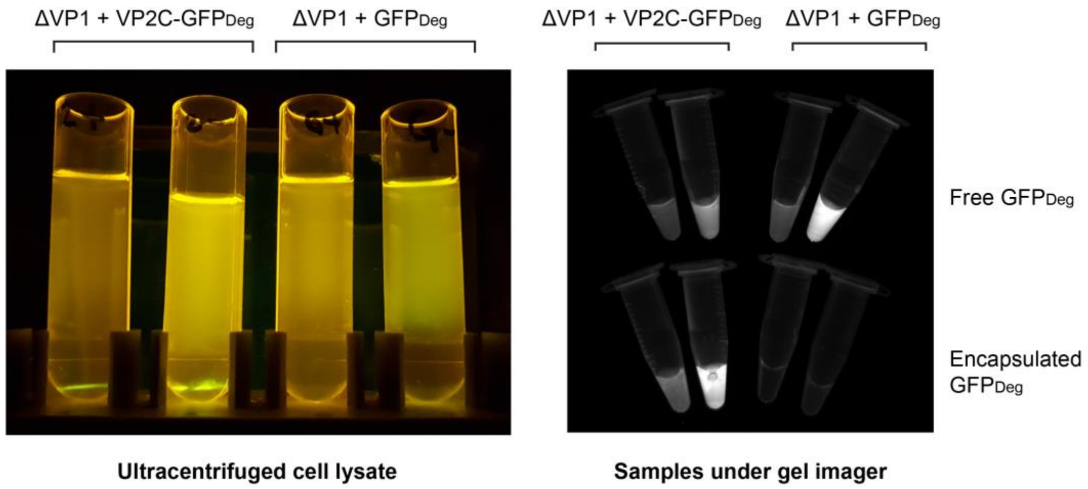

Figure S4. Flow cytometry to optimise the duration of cycloheximide treatment.

Data shows the 3 constructs from Figure 5 (ΔVP1 + VP2C-$GFP_{Deg}$, ΔVP1 + $GFP_{Deg}$, VP2C-$GFP_{Deg}$ only) along with controls expressing free destabilised GFP alone ($GFP_{Deg}$ only) and GFP without the degron tag (ΔVP1 + VP2C-GFP, ΔVP1 + GFP). Cultures were induced for 6 hours prior to adding cycloheximide (shown as time = 0). Each point shown is the mean of 3-4 biological replicates. $GFP_{Deg}$ strains with and without ΔVP1 showed very similar profiles, indicating that ΔVP1 does not affect GFP expression.

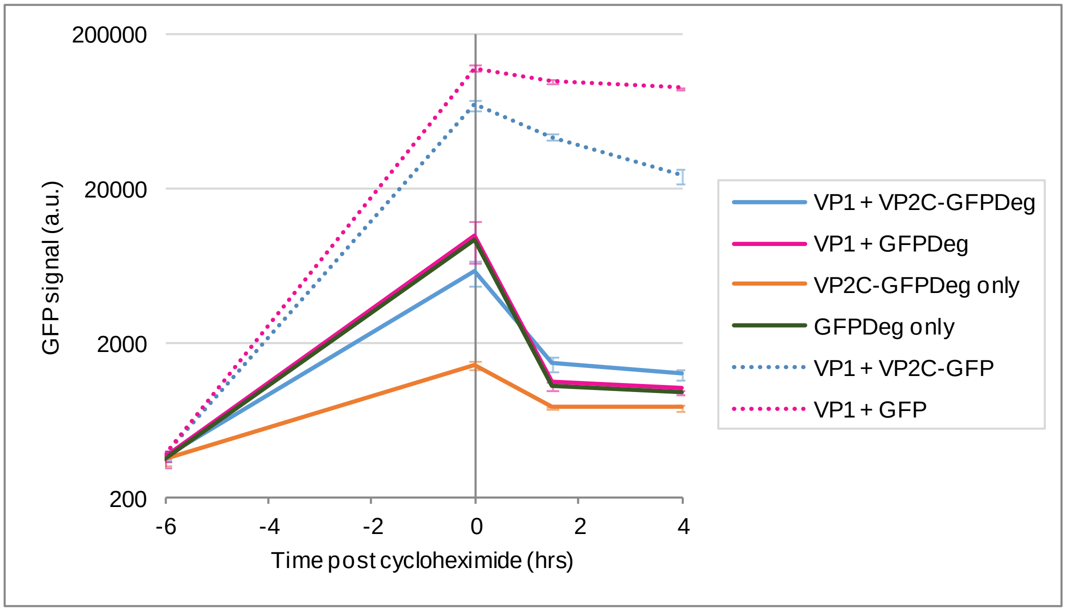

Figure S5. Confocal microscopy images of strains expressing VP2C-$GFP_{Deg}$ alone and a blank untransformed strain (negative control).

Image acquisition and display settings are the same as for Figure 4 in the main text.

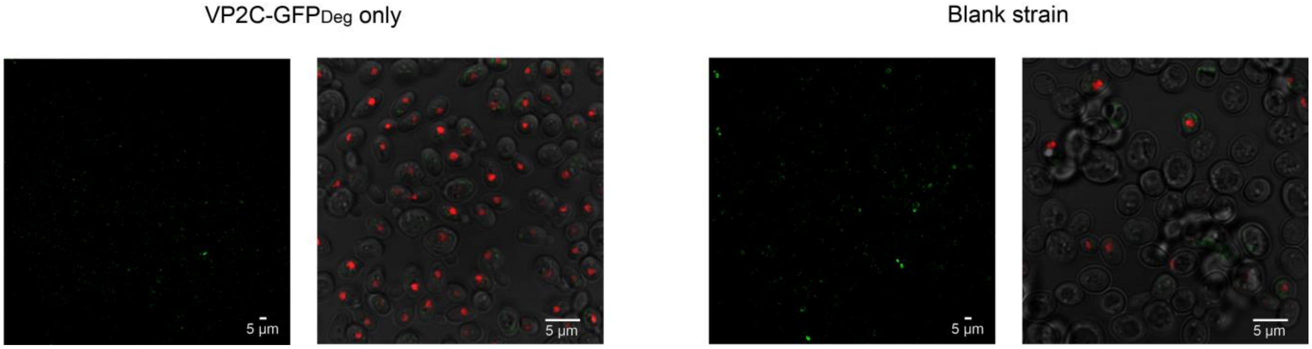

Figure S6. Examining the effect of wtVP1 and ΔVP1 expression of cell growth and protein expression.

Saturated YPD pre-cultures were inoculated into 5 ml YPG (in 28 ml McCartney bottles) at a starting OD600 of 0.2 and grown at 30 °C, 200 rpm shaking.

(a) Growth curves and (b) flow cytometry signal of strains with and without VP1. The graph legend for (b) is the same as in (a). Each point is the mean of 3 biological replicates, and error bars show the standard deviation.

(c) Anti-VP1 western blot of cultures sampled at 24, 48, and 72 hours post-induction. The same amount of sample was loaded in each lane, based on the $OD_{600}$ value. Similar expression levels of VP1 were observed for wtVP1 and ΔVP1, and VP1 levels were stable from 24-72 hours post-induction.

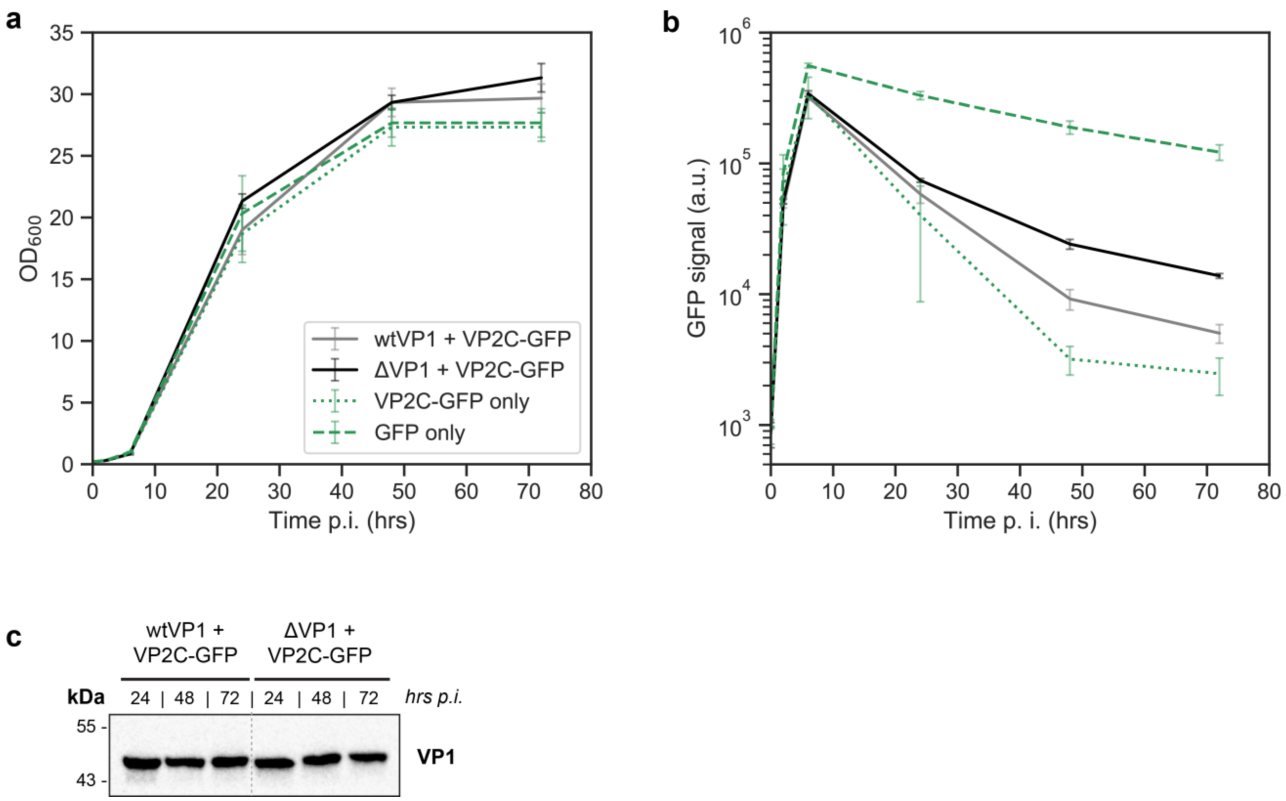

Figure S7. Compartmentalisation rescues growth defects from MIOX overexpression.

Growth profiles from Figure 7a in the main text are shown here overlaid with data from a non-MIOX-expressing control strain (ΔVP1 + VP2C-GFP). Strains were all grown simultaneously using the same medium and conditions. All data points are the means of 3 biological replicates; error bars are +/- 1 STD.

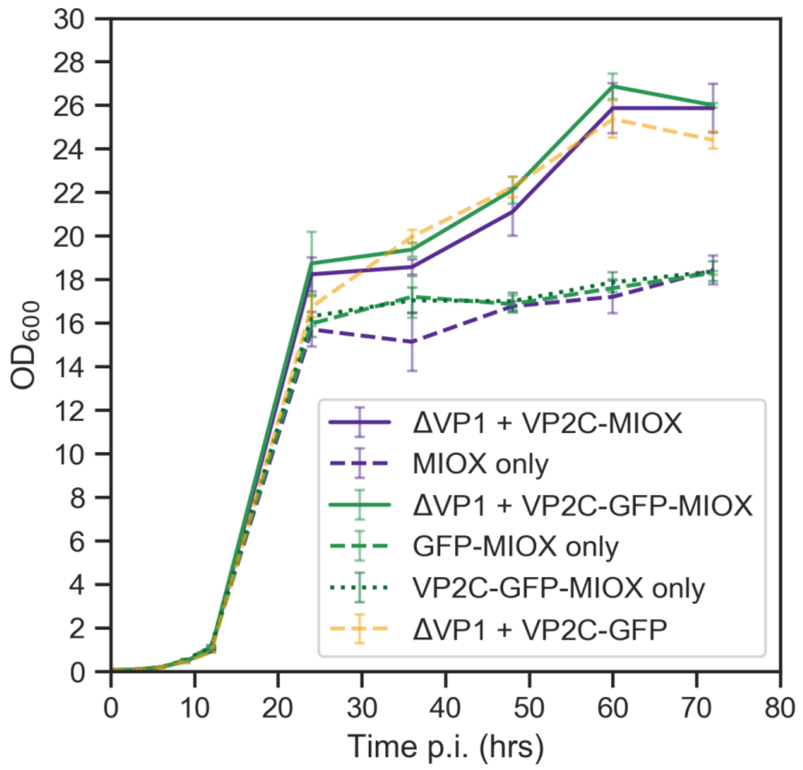

Table S1. PCR primer sequences.
All gene constructs were cloned by isothermal assembly. This method requires generating dsDNA fragments with a region at each end that overlaps with adjacent fragments.

| **Primer name** | **Description** | **Sequence (5' - 3')** | **Length (nt)** | **PCR template** | **PCR fragment generated** | **Used for which constructs?** | **Comments** |
|---|---|---|---|---|---|---|---|
| VP1_NheI_GAL1p_fw | ΔVP1 forward | CTTTAACGTCAAGGAGAAAAAACTATAGCTAGCAAAATGGCTTCAGGTGTAAG | 53 | Synthetic VP1 gene | ΔVP1 | ΔVP1 empty | VP1 replaces GFP sequence in pILGFPB5A |
| VP1_NotI_URA3t_rv | ΔVP1 reverse | CTTTGCAAATAGTCCTACTAGTGCGGCCGCTTAATTACCAGGGAATACAGTTTTAG | 56 | Synthetic VP1 gene | ΔVP1, wtVP1 | ΔVP1 empty, wtVP1 empty | |
| Fseq_GAL1p | Forward sequencing primer for VP1 | ACTTCTTATTCAAATGTCATAAAAGTATCAAC | 32 | NA | NA | ΔVP1 empty, wtVP1 empty | Binds to GAL1 promoter |
| Rseq_ScURA3t | Reverse sequencing primer for VP1 | CCAGCCTGCTTTTCTGTAAC | 20 | NA | NA | ΔVP1 empty, wtVP1 empty | Binds to *S. cerevisiae* URA3 terminator |
| Fseq_GAL10p | Forward sequencing primer for VP2C-cargo | GTGGTAATGCCATGTAATATGATTATTAAAC | 31 | NA | NA | All cargo constructs | Binds to GAL10 promoter |
| Rseq_KlURA3t | Reverse sequencing primer for VP2C-cargo | GCATTGGCACGGTGCAACAC | 20 | NA | NA | All cargo constructs | Binds to K. lactis URA3 terminator |
| NLS_fw_Gibson | wtVP1 forward; appends NLS residues to ΔVP1 | CGTCAAGGAGAAAAAACTATAGCTAGCAAAATGGCTCCAAAGAGAAAGTCAGGTGTAAGT | 72 | Synthetic VP1 gene | wtVP1 | wtVP1 empty, wtVP1 + VP2C-GFP, | wtVP1 replaces ΔVP1 |

| | | | | | | | |
|---|---|---|---|---|---|---|---|
| | | AAATGCGAAACC | | | | wtVP1 + GFP | |
| VP2_ClaI_GAL10p_fw | VP2C forward, with overhang for GAL10 promoter | AAAGTAAGAATTTTTGAAAATTCAATATAAATCGATAAAATGGGTAATGGTGGTCCAACT | 60 | Synthetic VP2C gene | VP2C | ΔVP1 + VP2C-GFP | VP2C-GFP-tHIS5 inserted between XbaI and EcoRI sites in VP1 plasmid |
| VP2_yEGFP_rv | VP2C linker reverse, with overhang for yeGFP | CTTCACCTTTAGAGGATCCCATAGAACCGCCGCCACCGGAGCC | 43 | Synthetic VP2C gene | VP2C | ΔVP1 + VP2C-GFP | |
| yEGFP_VP2_fw | yeGFP forward, with overhang for VP2C linker | GGCTCCGGTGGCGGCGGTTCTATGGGATCCTCTAAAGGTGAAG | 43 | pILGFPB5A (Peng et al. 2015[2]) | GFP for VP2C | ΔVP1 + VP2C-GFP | VP2C-GFP-tHIS5 inserted between XbaI and EcoRI sites in VP1 plasmid |
| yEGFP_HIS5t_rv | yeGFP reverse, with overhang for HIS5 terminator | ATAAAACTGTACATATACTGTTTAAATTAATCTATTTAAGATCTTTTGTACAATTCATCC | 60 | pILGFPB5A (Peng et al. 2015) | GFP for VP2C | ΔVP1 + VP2C-GFP | |
| HIS5t_yEGFP_fw | HIS5 terminator forward, with overhang for yeGFP | GGATGAATTGTACAAAAGATCTTAAATAGATTAATTTAAACAGTATATGTACAGTTTTAT | 60 | Yeast strain S288c genomic DNA | HIS5 terminator | ΔVP1 + VP2C-GFP | VP2C-GFP-tHIS5 inserted between XbaI and EcoRI sites in VP1 plasmid |
| HIS5t_KlURA3t_rv | HIS5 terminator reverse, with overhang for K. lactis URA3 terminator | TGATCTGTTGTATTGGGATCTCTAGAGTAACAATATCATGAGACCTTTTATAG | 53 | Yeast strain S288c genomic DNA | HIS5 terminator | ΔVP1 + VP2C-GFP | |
| yEGFP_ClaI_GAL10p_fw | yeGFP forward for controls without VP2C | AAAGTAAGAATTTTTGAAAATTCAATATAAATCGATAAAATGGGATCCTCTAAAGGTGA | 59 | pILGFPB5A (Peng et al. 2015) | Free GFP | ΔVP1 + GFP | GFP-tHIS5 inserted between XbaI and EcoRI sites in VP1 plasmid |

| | | | | | | | |
|---|---|---|---|---|---|---|---|
| GAL1p_patch | Re-circularises plasmid after VP1 excision | CTTTAACGTCAAGGAGAAAAAACTATAACTAGTAGGACTATTTGCAAAGG | 50 | NA | NA | VP2C-GFP only, VP2C-GFP-MIOX only | Oligomer used directly in isothermal assembly mix with plasmid; VP1 removed by digesting with NotI + NheI and gel purifying |
| ScLEU2t_fw | Forward primer for part of the LEU2 terminator | TGTCAAGGAAATCTTGGCTTAAAAAGATTCTCTTTTTTTATGATATTTGTACATAAACTT | 60 | Yeast strain S288c genomic DNA | LEU2 terminator fragment | UDH | Sequence inserted next to KlLEU2 in pUG73[3]; overlap region for leu2 gene replacement |
| ScLEU2t_SwaI_rv | Reverse primer for part of the LEU2 terminator | GCCACCTGACGTCATTTAAATCAATATTAATGTTAAAGTGCAATTCTTTTTCCTTATCAC | 60 | Yeast strain S288c genomic DNA | LEU2 terminator fragment | UDH | |
| ScLEU2p_SwaI_fw | Forward primer for part of the LEU2 promoter | GTGAGCGAGGAAGCATTTAAATAACTGTGGGAATACTCAGGTATCGTAAGATG | 53 | Yeast strain S288c genomic DNA | LEU2 promoter fragment | UDH | Sequence inserted next to tCYC1; overlap region for leu2 gene replacement |
| ScLEU2p_NotI_rv | Reverse primer for part of the LEU2 promoter | CACTAGTGGCCTATGCGGCCGCTAGAATGGTATATCCTTGAAATATATATATATATATTG | 60 | Yeast strain S288c genomic DNA | LEU2 promoter fragment | UDH | |
| pTEF1_EcoRI_fw | TEF1 promoter forward | TGGAAATCATTGAGTCGATGAATTCCCACACACCATAGCTTC | 42 | pILGFPE9 (Peng et al. 2015[2]) | TEF1 promoter | UDH | pTEF1, UDH, and tCYC1 co-assembled with |

| | | | | | | | |
|---|---|---|---|---|---|---|---|
| pTEF1_XhoI_rv | TEF1 promoter reverse | TTTCTCGAGTTTGTAATTAAAACTTA | 26 | pILGFPE9 (Peng et al. 2015) | TEF1 promoter | UDH | linearised backbone vector |
| UDH_pTEF1_fw | UDH forward, with overhang for TEF1 promoter | CTAAGTTTTAATTACAAACTCG | 22 | Synthetic UDH gene | UDH | UDH | pTEF1, UDH, and tCYC1 co-assembled with linearised backbone vector |
| UDH_SbfI_rv | UDH reverse, with overhang for CYC1 terminator | GATGCGGCCCTCCTGCAGGTTATTTATCGCCGAACGGTCC | 40 | Synthetic UDH gene | UDH | UDH | |
| tCYC1_SbfI_fw | CYC1 terminator forward, with overhang for UDH | CCTGCAGGAGGGCCGCATCAT | 21 | pCM251 (Belli et al. 1998[4]) | CYC1 terminator | UDH | pTEF1, UDH, and tCYC1 co-assembled with linearised backbone vector |
| tCYC1_NotI_rv | CYC1 terminator reverse, with overhang for LEU2 promoter | CAAGGATATACCATTCTAGCGGCCGCAGGCCGCAAATTAAAGC | 43 | pCM251 (Belli et al. 1998) | CYC1 terminator | UDH | |
| Fseq_TEF1p | Forward sequencing primer for UDH | TTTACTTCTTGCTCATTAGAAAGAAAG | 27 | NA | NA | UDH | Binds to TEF1 promoter |
| Rseq_CYC1t | Reverse sequencing primer for UDH | TTGTCTAACTCCTTCCTTTTCGGTTAGAG | 29 | NA | NA | UDH | Binds to CYC1 terminator |
| yEGFP_AG4TGGA_rv | yeGFP reverse for GFP-MIOX fusion proteins; appends a linker peptide (AGGGGTGGAEL) sequence | AGCACCACCAGTACCTCCTCCACCAGCAGATCTTTTGTACAATTCATCCATACCATG | 57 | pILGFPB5A (Peng et al. 2015) | GFP for GFP-MIOX fusions | VP2C-GFP-MIOX + VP1, VP2C-GFP-MIOX only, GFP-MIOX only | Fragment co-assembled with mmMIOX gene fragment and linearised backbone vector |

Table S2. Synthetic gene sequences.

| Synthetic dsDNA | Description | Sequence (5' - 3') | Length (nt) | Comments |
|---|---|---|---|---|
| ΔVP1 (in pUC57 vector) | Murine polyomavirus VP1, NLS removed (ΔVP1 mutant), codon-optimised for *S. cerevisiae* | ATGGCTTCAGGTGTAAGTAAATGCGAAACCAAATGCACAAAGGCTTGCCCT<br>AGACCAGCTCCAGTTCCTAAGTTATTGATTAAAGGTGGTATGGAAGTTTTGG<br>ATTTGGTTACAGGTCCAGATTCTGTTACTGAAATCGAAGCATTTTTGAACCC<br>AAGAATGGGTCAACCACCAACACCAGAATCTTTAACTGAAGGTGGTCAATA<br>TTACGGTTGGTCAAGAGGTATTAATTTGGCTACATCTGATACTGAAGATTCA<br>CCAGGTAATAATACATTACCAACTTGGTCTATGGCAAAATTGCAATTACCAA<br>TGTTGAACGAAGATTTGACTTGTGATACTTTGCAAATGTGGGAAGCAGTTTC<br>AGTTAAAACAGAAGTTGTTGGTTCTGGTTCTTTGTTGGATGTTCATGGTTTTA<br>ATAAGCCAACAGATACTGTTAACACAAAGGGTATTTCTACTCCAGTTGAAGG<br>TTCACAATATCATGTTTTTGCTGTTGGTGGTGAACCATTGGATTTGCAAGGTT<br>TAGTTACAGATGCAAGAACTAAGTACAAGGAAGAAGGTGTTGTTACTATTA<br>AAACAATCACTAAGAAAGATATGGTTAATAAGGATCAAGTTTTGAACCCAA<br>TTTCAAAGGCTAAATTGGATAAGGATGGCATGTACCCAGTTGAAATTTGGCA<br>TCCAGATCCAGCTAAAAATGAAAACACAAGATACTTCGGTAATTACACTGG<br>TGGTACTACAACTCCACCAGTTTTGCAATTCACTAACACTTTGACAACTGTTT<br>TGTTAGATGAAAATGGTGTTGGTCCATTATGTAAAGGTGAAGGTTTGTACTT<br>ATCTTGCGTTGATATTATGGGTTGGAGAGTTACAAGAAACTACGATGTTCAT<br>CATTGGAGAGGTTTGCCAAGATACTTCAAGATTACTTTAAGAAAAAGATGG<br>GTTAAAAATCCATACCCAATGGCTTCTTTGATTTCTTCTTTGTTTAATAATAT<br>GTTACCACAAGTTCAAGGTCAACCAATGGAAGGTGAAAACACACAAGTTGA<br>AGAAGTTAGAGTTTACGATGGTACTGAACCAGTTCCAGGTGACCCTGACATG<br>ACAAGATATGTAGATAGATTCGGTAAAACTAAAACTGTATTCCCTGGTAATT<br>AA | 1143 | PCR template |
| VP2C_linker_yco | Murine polyomavirus VP2(251-301)- | ATGGGTAATGGTGGTCCAACTCCAGCTGCTCATATTCAAGATGAATCTGGTG<br>AAGTTATTAAATTTTATCAAGCTCAAGTTGTTTCTCATCAAAGAGTTACTCCA<br>GATTGGATGTTGCCATTGATTTTGGGTTTGTATGGTGACATCACTCCAGGTG<br>GTGGTGGTTCTGGTGGCGGTGGCTCCGGTGGCGGCGGTTCT | 198 | PCR template |

| | | | | |
|---|---|---|---|---|
| | [GGGGS]3, codon-optimised for *S. cerevisiae* | | | |
| mODC(425-461) | Mouse ornithine decarboxylase fragment (degron region), codon-optimised for *S. cerevisiae* | GCTATGGTATGGATGAATTGTACAAAAGATCTTTTCCTCCTGAAGTTGAAGA<br>GCAAGATGATGGAACTTTGCCAATGTCATGTGCACAAGAATCAGGTATGGA<br>TAGGCATCCAGCTGCTTGTGCATCTGCAAGAATTAATGTTTAAATAGATTAA<br>TTTAAACAGTATATGTACAG | 175 | Fragment used directly in isothermal assembly mix with BglII-digested plasmid |
| psUDH | *P. syringae* UDH, codon-optimised for *S. cerevisiae*; fragment synthesised with assembly overlaps | GCCCTCCTGCAGGTTATTTATCGCCGAACGGTCCGGACGCTACAAATGCACC<br>GCCCTGGTAAACCATTGCAGGGTCGTCGTCAGCTGGCATTGGCTGGGCGTCG<br>ACCTTGGCACGGAAGACCTCCGAGCTGTCTTTAGGGGCGTAGTCCAGTTTGC<br>TGGCAAATCTGTTGTCCCACCAAACGGTCTTGTTGTCAGACACGCCATACAC<br>CACGGTGTGGCCGACGTCCGGCGTGTACAGGGCACGCTCTAGCAAACGGGT<br>CAGGTCATCGAAGCTCAGCCAGGTGCTCATCATTCTACGGTTCTGTGGTTCA<br>GGGAAGGATGAGCCGATACGGATGCTGACGGTTTCGATGCCGTAACGATCA<br>AAGTAGAAGCTGGCCATGTCTTCGCCGTAGGACTTGGACAAACCATAGTAA<br>GAATCTGGCCTACGTGGGGAGTGCGCGTCGATGGTTTCGTTCTGCTTGTAGA<br>AACCGATAACATGGTTGGAGCTGGCGAAGATCACACGTTTCACGCCATGAC<br>GACGGGCGGCCTCGTAGATATGGAAAACGCCGCAGATATTAGCGCCCAAGA<br>TTTCCTCGAAAGGTCTTTCGACAGACACGCCGCCAAAATGCAAAATGGCGTC<br>AACGCCTTCGACCAGACGGTGTACGGCGTCTTTGTCCGCCAGATCGCAGACC<br>TGAACTTCTTCATGGTCGCCGACAGCGGGAGCCATCTCGGCGATATCAGATA<br>GACGCAGAATGTGTGAGTAAGGACGCAATGTCTCACGCAATACCTTGCCCA<br>GGCCGCCTGCAGCACCGGTCAGCAGTAGACGATTGAATGGAGTTTGAGTAG<br>TATGAGCAGATGCCATTTTCTCGAGTTTGTAATTAAAACTTAG | 868 | PCR template |

| | | | | |
|---|---|---|---|---|
| mmMIOX | Mouse (*Mus musculus*) MIOX, codon-optimised for *S. cerevisiae*; fragment synthesised with assembly overlaps | AGGTACTGGTGGTGCTGAGCTCATGAAGGTCGATGTTGGTCCAGATCCTTCC CTGGTCTATCGTCCCGATGTGGACCCAGAGATGGCCAAAAGCAAGGACAGC TTCCGTAACTATACTTCAGGCCCGTTGCTGGATCGTGTCTTTACCACATACA AGCTGATGCACACTCACCAGACTGTGGACTTCGTCAGCAGGAAGCGTATCC AGTATGGAGGCTTCTCTTACAAGAAGATGACCATCATGGAGGCTGTGGGCA TGCTGGATGATCTGGTGGACGAATCTGACCCAGACGTAGATTTCCCCAACTC CTTCCACGCGTTCCAGACCGCGGAGGGCATCCGTAAAGCCCACCCGGACAA GGACTGGTTCCACCTGGTCGGACTTTTGCACGATCTGGGGAAAATTATGGCT CTGTGGGGGGAACCTCAGTGGGCTGTTGTTGGAGACACGTTCCCCGTGGGCT GCCGTCCCCAGGCCTCTGTGGTGTTCTGTGACTCTACTTTCCAGGACAATCCT GACCTGCAGGATCCTCGTTACAGCACAGAACTGGGCATGTACCAGCCTCACT GTGGACTAGAGAACGTCCTTATGTCCTGGGGCCATGATGAGTACCTATACCA GATGATGAAGTTCAACAAGTTCTCCCTGCCTTCAGAGGCCTTCTACATGATC CGTTTCCACTCCTTCTATCCGTGGCACACCGGCGGTGACTACCGTCAGCTGT GCAGCCAGCAGGACCTGGATATGCTGCCCTGGGTGCAAGAGTTCAACAAGT TTGATCTGTACACGAAGTGCCCTGACCTACCGGATGTGGAGAGCCTGCGTCC CTACTATCAAGGGCTGATTGACAAGTACTGCCCGGGCACCCTGAGCTGGTG ACTCGAGATAGATTAATTTAAACAGTATATGTACAGTTTTA | 920 | Fragment used directly in isothermal assembly mix with linearised backbone; yeGFP first removed from backbone by digesting with BamHI + BglII and gel purifying |

Table S3. Yeast strain, plasmid, and protein details.

Except for D-glucaric acid expression strains, every yeast strain in this work was CEN.PK2-1C transformed with the corresponding URA3 YIp. For D-glucaric acid expression, CEN.PK2-1C was first transformed with the UDH (LEU2) YIp; the same recovered colony was then transformed with the corresponding MIOX cassettes (URA3 YIp). Transformation of each plasmid results in single copy integration of the expression cassette in the genome.

| **Strain** | **Description** | **Reference** |
|---|---|---|
| CEN.PK2-1C | Blank base strain;<br>MATa, his3D1, leu2-3_112, ura3-52, trp1-289, MAL2-8c, SUC2 | Entian & Kötter 2007[5] |
| **Plasmid** | **Description** | **Reference** |
| pILGFPB5A | Backbone vector; pGAL1-pGAL10-yeGFP, URA3, YIp | Peng et al. 2015[2] |
| wtVP1 only | pGAL1-wtVP1-tURA3, URA3, YIp | This work |
| wtVP1 + VP2C-GFP | pGAL1-wtVP1-tURA3 + pGAL10-VP2C-yeGFP-tHIS5, URA3, YIp | This work |
| ΔVP1 only | pGAL1-ΔVP1-tURA3, URA3, YIp | This work |
| ΔVP1 + VP2C-GFP | pGAL1-ΔVP1-tURA3 + pGAL10-VP2C-yeGFP-tHIS5, URA3, YIp | This work |
| UDH only | pTEF1-UDH-tCYC1, LEU2, YIp | This work |
| ΔVP1 + VP2C-MIOX | pGAL1-ΔVP1-tURA3 + pGAL10-VP2C-MIOX-tHIS5, URA3, YIp | This work |
| MIOX only | pGAL10-MIOX-tHIS5, URA3, YIp | This work |
| ΔVP1 +<br>VP2C-GFP-MIOX | pGAL1-ΔVP1-tURA3 + pGAL10-VP2C-yeGFP-MIOX-tHIS5, URA3, YIp | This work |
| GFP-MIOX only | pGAL10-yeGFP-MIOX-tHIS5, URA3, YIp | This work |
| VP2C-GFP-MIOX only | pGAL10-VP2C-yeGFP-MIOX-tHIS5, URA3, YIp | This work |

| **Protein** | **Sequence** | **Reference** |
|---|---|---|
| wtVP1 | MAPKRKSGVSKCETKCTKACPRPAPVPKLLIKGGMEVLDLVTGPDSVTEIEAFLNPRMGQPPT<br>PESLTEGGQYYGWSRGINLATSDTEDSPGNNTLPTWSMAKLQLPMLNEDLTCDTLQMWEAV<br>SVKTEVVGSGSLLDVHGFNKPTDTVNTKGISTPVEGSQYHVFAVGGEPLDLQGLVTDARTKY<br>KEEGVVTIKTITKKDMVNKDQVLNPISKAKLDKDGMYPVEIWHPDPAKNENTRYFGNYTGGT<br>TTPPVLQFTNTLTTVLLDENGVGPLCKGEGLYLSCVDIMGWRVTRNYDVHHWRGLPRYFKIT<br>LRKRWVKNPYPMASLISSLFNNMLPQVQGQPMEGENTQVEEVRVYDGTEPVPGDPDMTRYV<br>DRFGKTKTVFPGN | UniProtKB -<br>P49302<br>(VP1_POVMP) |
| ΔVP1 | MASGVSKCETKCTKACPRPAPVPKLLIKGGMEVLDLVTGPDSVTEIEAFLNPRMGQPPTPESLT<br>EGGQYYGWSRGINLATSDTEDSPGNNTLPTWSMAKLQLPMLNEDLTCDTLQMWEAVSVKTE<br>VVGSGSLLDVHGFNKPTDTVNTKGISTPVEGSQYHVFAVGGEPLDLQGLVTDARTKYKEEGV<br>VTIKTITKKDMVNKDQVLNPISKAKLDKDGMYPVEIWHPDPAKNENTRYFGNYTGGTTTPPV<br>LQFTNTLTTVLLDENGVGPLCKGEGLYLSCVDIMGWRVTRNYDVHHWRGLPRYFKITLRKR<br>WVKNPYPMASLISSLFNNMLPQVQGQPMEGENTQVEEVRVYDGTEPVPGDPDMTRYVDRFG<br>KTKTVFPGN | Catrice & Sainsbury<br>2005[6] |
| VP2C-GFP<br>==<br>VP2C-[GGGGS]$_3$-<br>yeGFP<br><br>VP2C == VP2(251-<br>301) | MGNGGPTPAAHIQDESGEVIKFYQAQVVSHQRVTPDWMLPLILGLYGDITPGGGGSGGGGSG<br>GGGSMGSSKGEELFTGVVPILVELDGDVNGHKFSVSGEGEGDATYGKLTLKFICTTGKLPVPW<br>PTLVTTFGYGVQCFARYPDHMKQHDFFKSAMPEGYVQERTIFFKDDGNYKTRAEVKFEGDTL<br>VNRIELKGIDFKEDGNILGHKLEYNYNSHNVYIMADKQKNGIKVNFKIRHNIEDGSVQLADHY<br>QQNTPIGDGPVLLPDNHYLSTQSALSKDPNEKRDHMVLLEFVTAAGITHGMDELYKRS | VP2C references:<br>Catrice & Sainsbury<br>2005[6]<br>UniProtKB -<br>P12908<br>(VP2_POVMC)<br><br>yeGFP reference:<br>Cormack et al.<br>1997[7] |
| VP2C-GFP$_{Deg}$<br>==<br>VP2C-[GGGGS]$_3$-<br>yeGFP- mODC(425-<br>461) | MGSSKGEELFTGVVPILVELDGDVNGHKFSVSGEGEGDATYGKLTLKFICTTGKLPVPWPTLV<br>TTFGYGVQCFARYPDHMKQHDFFKSAMPEGYVQERTIFFKDDGNYKTRAEVKFEGDTLVNRI<br>ELKGIDFKEDGNILGHKLEYNYNSHNVYIMADKQKNGIKVNFKIRHNIEDGSVQLADHYQQN<br>TPIGDGPVLLPDNHYLSTQSALSKDPNEKRDHMVLLEFVTAAGITHGMDELYKRSFPPEVEEQ<br>DDGTLPMSCAQESGMDRHPAACASARINV | Hoyt et al. 2003[8]<br><br>mODC sequence:<br>UniProtKB -<br>P00860 |

| | | |
|---|---|---|
| | | (DCOR_MOUSE) |
| *M. musculus* MIOX | MKVDVGPDPSLVYRPDVDPEMAKSKDSFRNYTSGPLLDRVFTTYKLMHTHQTVDFVSRKRIQYGGFSYKKMTIMEAVGMLDDLVDESDPDVDFPNSFHAFQTAEGIRKAHPDKDWFHLVGLLHDLGKIMALWGEPQWAVVGDTFPVGCRPQASVVFCDSTFQDNPDLQDPRYSTELGMYQPHCGLENVLMSWGHDEYLYQMMKFNKFSLPSEAFYMIRFHSFYPWHTGGDYRQLCSQQDLDMLPWVQEFNKFDLYTKCPDLPDVESLRPYYQGLIDKYCPGTLSW | UniProtKB - Q9QXN5 (MIOX_MOUSE) |
| *P. syringae* UDH | MASAHTTQTPFNRLLLTGAAGGLGKVLRETLRPYSHILRLSDIAEMAPAVGDHEEVQVCDLADKDAVHRLVEGVDAILHFGGVSVERPFEEILGANICGVFHIYEAARRHGVKRVIFASSNHVIGFYKQNETIDAHSPRRPDSYYGLSKSYGEDMASFYFDRYGIETVSIRIGSSFPEPQNRRMMSTWLSFDDLTRLLERALYTPDVGHTVVYGVSDNKTVWWDNRFASKLDYAPKDSSEVFRAKVDAQPMPADDDPAMVYQGGAFVASGPFGDK | UniProtKB - Q888H1 (URODH_PSESM) |

Figure S8. SEC polishing step for VLP purification.

(a) During iodixanol cushion ultracentrifugation, MPyV VLP co-sediments with two high-MW yeast contaminants that are visible under negatively-stained TEM: the fatty acid synthase complex (FAS), and a ubiquitous yeast virus Saccharomyces cerevisiae virus LA (ScV-LA).

(b) Sample SDS-PAGE of ultracentrifuge-purified MPyV showing the major contaminants. Protein identity was determined by LC-MS/MS of excised bands.

(c) A sample SEC run profile (HiPrep 16/60 Sephacryl S-500 HR, GE Healthcare). Iodixanol absorbs very strongly at A280 and is seen as a massive peak (>2000 mAU) near the end of 1 CV (120 ml). Fractions eluting between 50-70 ml (indicated by dotted grey lines) were pooled and filter concentrated for further analysis.

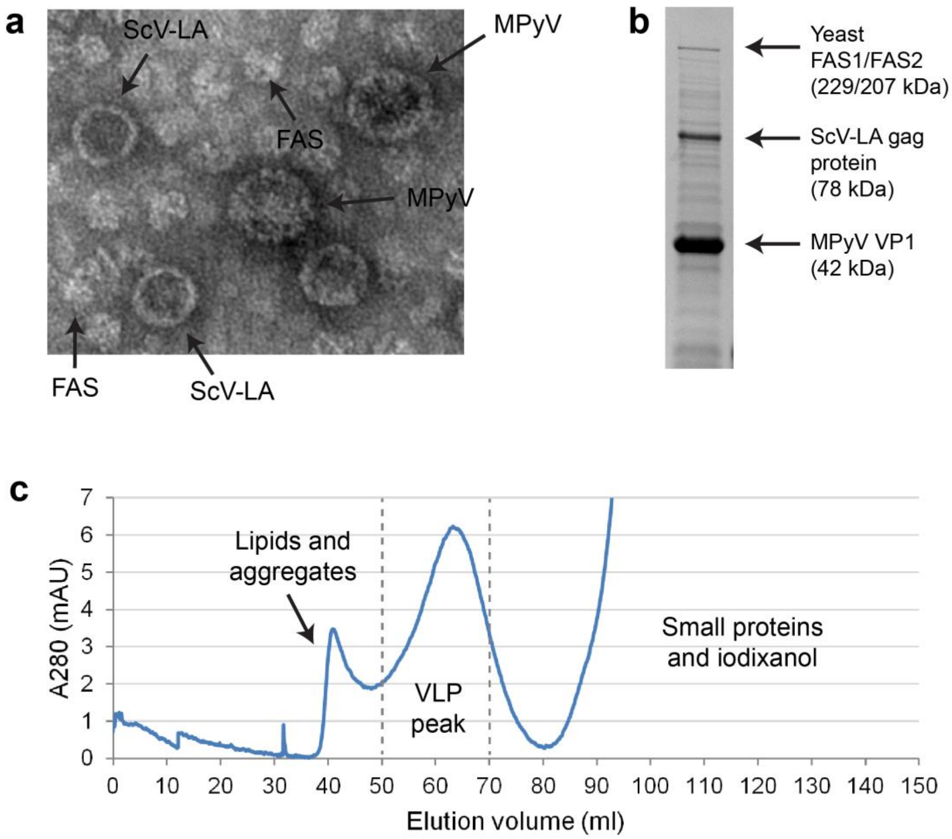
